# Spatiotemporal encoding of touch signals in the human somatosensory and motor cortices

**DOI:** 10.64898/2026.05.08.721831

**Authors:** Michela Cattabriga, Ali H. Alamri, Taylor G. Hobbs, Alexandriya M. X. Emonds, Anton R. Sobinov, Robert A. Gaunt, Charles M. Greenspon, Giacomo Valle

## Abstract

The sense of touch is fundamental for dexterous manipulation, object interaction, and body awareness. It is primarily processed in the somatosensory cortex (SC), yet our understanding of how tactile information is encoded at the level of neural populations and single neurons in humans remains limited. It is unclear how natural tactile signals are represented in SC and how they may be influenced by visual inputs, as well as how closely sensory and motor cortices interact during passive touch. Here, we investigated the neural basis of touch in the human SC using chronically implanted microelectrode arrays in three participants. By delivering controlled mechanical stimuli, we characterized neural responses to natural touch and mapped detailed somatotopic receptive fields (the patch of skin that elicits neural responses when stimulated) in humans, including multi-digit representations. Surprisingly, we also found strong, clearly somatotopic activation in the motor cortex (MC) during passive touch, even in the absence of movement, highlighting a tight and functionally relevant sensorimotor coupling. We further examined how vision shapes tactile processing by comparing neural activity during actual touch with and without vision, and during observation of touch on another person’s hand. While touch to the participants’ hands elicited robust, event-locked, and somatotopically organized responses in the SC, observation of tactile actions alone did not produce significant activation, suggesting limited vicarious encoding at this level. These findings provide a detailed characterization of human touch processing at the level of neuronal populations and give insights for the design of microstimulation strategies of the SC for the restoration of touch.

## Introduction

Human intelligence has evolved in tandem with the hand, our principal means to interact with the world. The sense of touch plays a crucial role in the human manual dexterity^1^ enabling precise object manipulation, grip force modulation, texture discrimination, and real-time feedback necessary for coordinated and adaptive movements^2^. Indeed, the absence of tactile feedback significantly impairs the ability to grasp and manipulate objects^3^. Tactile afferent signals from the periphery ascend along the neuroaxis via the dorsal column–medial leminiscal and spinothalamic pathways, undergoing successive processing in the spinal cord, brainstem nuclei, and thalamus (primarily the ventral posterior nuclei), before reaching the somatosensory cortex (SC)^4–6^. Within the SC and interconnected cortical areas, these signals are further integrated and transformed, contributing to the distributed neural processes underlying conscious somatosensory perception^4^. The functional organization, somatotopy, and response properties of the cortical neurons in the SC during corporeal touch have been extensively described in non-human primates (NHPs)^7–10^ and, mainly using non-invasive techniques, in humans^11,12^. However, the topographic organization and response properties of neurons within the SC during tactile stimulation in humans remain only partially characterized, especially at the neuronal level. In addition, previous animal studies have demonstrated that somatosensory signals can also reach the primary motor cortex (M1) through both thalamocortical pathways^13–15^ and cortico-cortical projections coming from SC to motor cortex (MC)^16–18^. Nevertheless, the precise mechanisms and cortical representation underlying the sensorimotor interplay remains poorly understood, partly due to the limited availability of high-channel-count neural recording systems capable of simultaneously sampling activity from both the SC and MC in humans.

Finally, the extent to which SC integrates visuo-tactile information and whether it encodes tactile-related events that are only visually observed is still unclear. Indeed, multiple fMRI studies in humans have reported activation in somatosensory cortical areas during the observation of touch applied to another person’s body^19^ (vicarious or mirror-like activity). In particular, observing touch delivered to another individual’s fingers, hands, or legs can activate the corresponding regions in the observer’s primary or second SC^20–22^. Additionally, SC activity can be enhanced when tactile stimulation is paired with visual information, but primarily when touch is applied to one’s own body rather than to someone else’s^23^. This evidence raises the possibility that multisensory integration may occur at the early stages of somatosensory processing and may generate vicarious responses in SC. However, other studies have failed to observe such effects during early stages of tactile processing^23–26^, leaving the question open of whether SC directly encodes observed tactile information.

To address all these questions, we simultaneously recorded neural population activity from the SC (Broadman area 1, BA1) and the MC (Broadman area 4 and area 6 dorsal, BA4 – BA6d) in three human participants in response to stimulation of the hand during different visuo-tactile paradigms. These participants had suffered a cervical spinal cord injury (SCI) yet retained partial tactile sensation. They were implanted with multiple microelectrode arrays in the SC and MC, providing a unique opportunity to investigate touch at the single-neuron level in humans, as well as how somatosensory and motor cortical populations encode touch, and how these regions interact during tactile processing.

## Results

Neural activity evoked by mechanical tactile stroking of individual digits was simultaneously recorded from the SC and MC of three participants with SCI using four intracortical microelectrode arrays (**Fig. 1A**). Two 96-channel Neuroport electrodes (Blackrock Neurotech, USA) were implanted in the arm and hand representation in MC, between BA6d and BA4, and two 32-channel Neuroport electrodes (Blackrock Neurotech, USA) in the hand representation in SC, BA1^27,28^ (**Fig. 1A**). Population responses, including both spatial and temporal neural features, were recorded and analyzed across different digits (**Fig. 1 B)** and three experimental conditions (**Fig. 1 C**): *touch with vision, touch without vision, and observed touch*. These intracortical recordings allowed us to investigate how SC and MC encode both actual and observed touch, as well as their potential interaction during tactile processing.

**Fig. 1.**
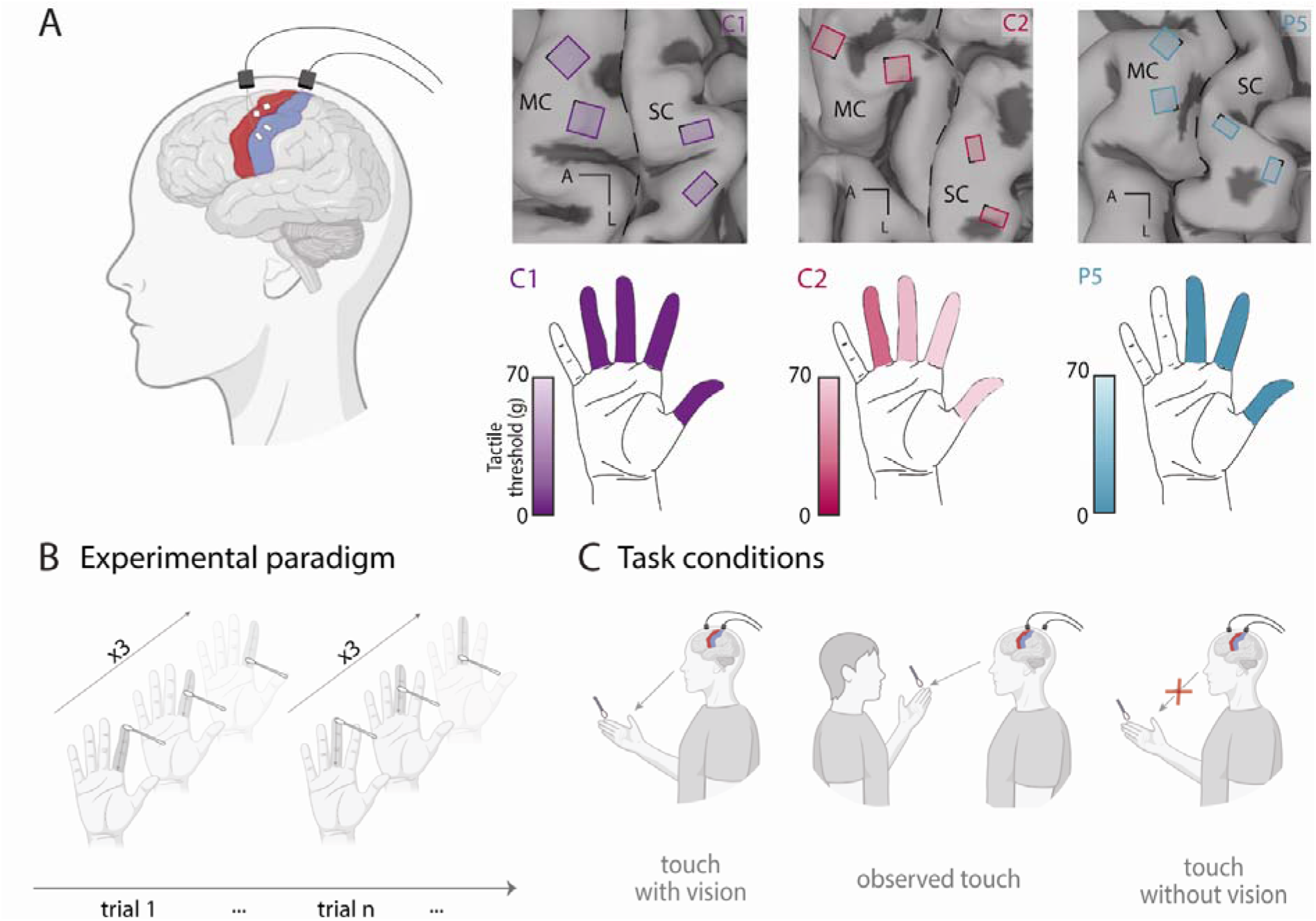
Investigating neural encoding of touch via intracortical interfacing. **(A)** Array implant locations overlaid on anatomical MRI scans. Four intracortical microelectrode arrays (Blackrock Neurotech, Inc.) were implanted in the hand and arm representations of the motor cortex (MC) and in the hand representation of the somatosensory cortex (SC, Brodmann Area 1). Implant locations are shown for participants C1, C2, and P5. Black dashed lines indicate the central sulcus. L, lateral; A, anterior. Bottom panel illustrates tactile thresholds normalized across participants on the stimulated fingers. Darker colors indicate lower tactile thresholds, and lighter colors indicate higher tactile thresholds. **(B)** Experimental paradigm. Each trial consisted of three consecutive tactile stimulations delivered to the same digit. **(C)** Task conditions. Three experimental conditions were tested: touch with vision, observed touch, and touch without vision.

### Cortical neurons in BA1 show both single and multi-digit tuning

We first characterized multi-unit activity in BA1 during actual touch with visual feedback to examine the tuning properties of the recorded neurons. Tuning was assessed by analyzing firing rates aligned to touch onset. We calculated firing rates from single channels responding to stimulation of the digits across the three participants (**Fig. 2A**). Across participants, 44% of the electrodes in the SC exhibited tuning to at least one digit, showing significant modulation in response to tactile events (C1: 69%; C2: 19%; P5: 44%; **Fig. 2B**). Among these responsive electrodes, 46% responded to stimulation of more than one digit (C1: 55%; C2: 33%; P5: 39%; **Fig. 2C**), indicating the presence of multi-digit RFs in BA1 (**Fig. 2A**, C1). Within the multi-digit RFs, most electrodes responded to two digits (**Fig. 2D**). Additionally, we assessed whether multi-digit electrodes maintained similar firing rates across digits or exhibited digit selectivity with rate coding by computing the absolute tolerance index. The results showed limited variation in firing rates across digits, with similar peak responses across digits (absolute tolerance index: C1=0.77±0.18, C2=0.86±0.07, P5=0.86 ±0.09) (**Fig. 2E**). Given the relative placement of the arrays and the fingers where the stimuli were applied, across digits, the index (D2) and middle (D3) were the most represented (**Fig. 2F**). Finally, touch-evoked responses of the cortical neurons in SC were predominantly excitatory, characterized by an increase in firing rate (**Fig. 2G**). To further characterize the spatial organization of these responses, we computed RF maps for each participant considering only the modulating electrodes. The resulting maps revealed the canonical somatotopic organization of BA1, with adjacent digits represented in continuous cortical regions (**Fig. 2H; Supp. Fig. 1-2**). Specifically, digit representations progressed from thumb (D1) to ring (D4) from the lateral to the medial array axis.

**Fig. 2.**
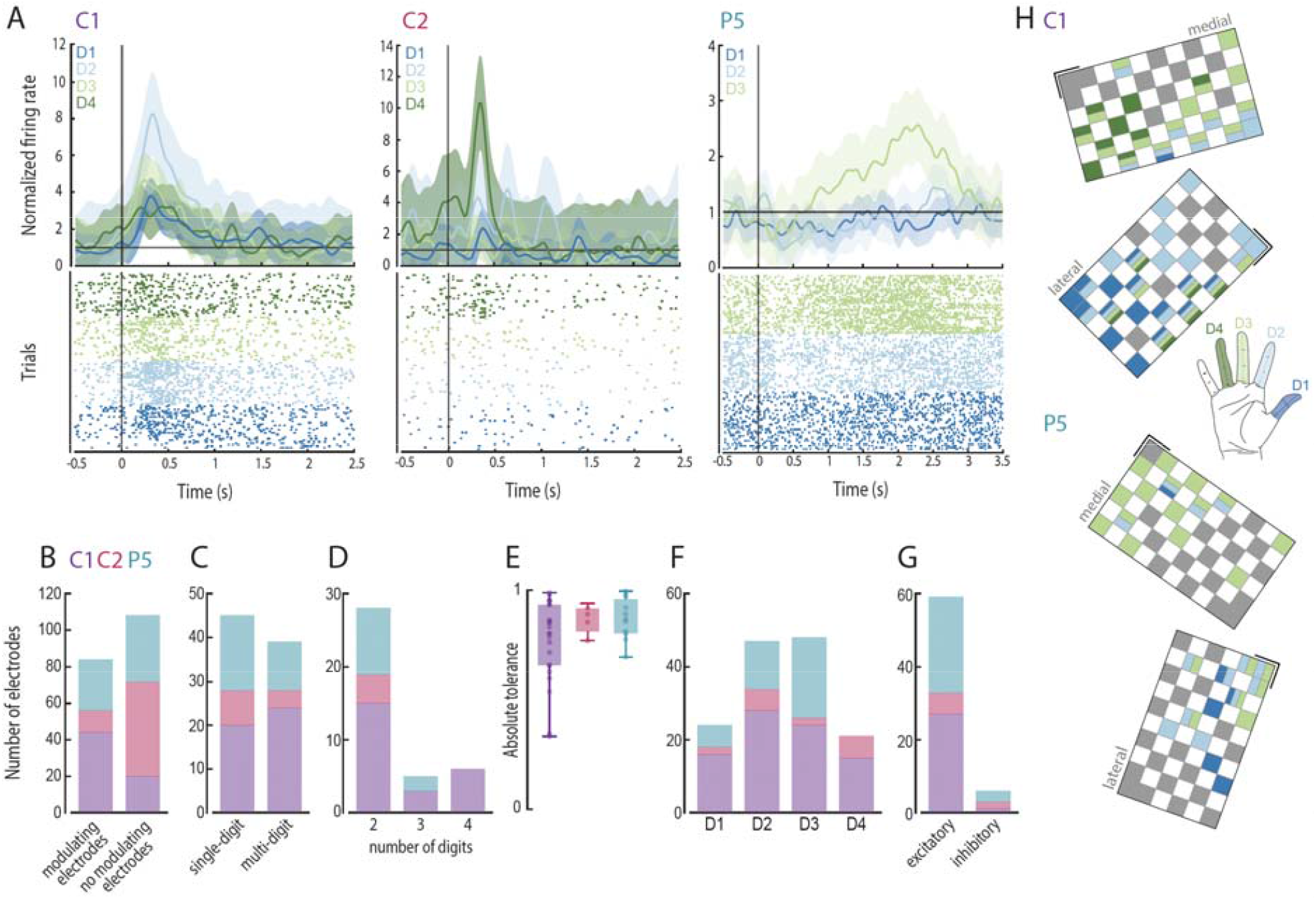
Touch-evoked responses in the human somatosensory cortex (SC). **(A)** Example of smoothed normalized firing rates and raster plots from sensory array channels in each participant in response to stimulation of each digit. Shaded areas indicate standard deviation. The plot on the left illustrates multi-digit activation and the plots on the middle and right show single-digit activation. Black vertical line indicates touch onset. **(B-D)** Bar plots summarizing the total number of modulating electrodes (n=192) **(B)**, the number of single-digit and multi-digit tuned electrodes (n=84) **(C)**, the distribution of the number of digits represented by multi-digit electrodes (n=39) **(D). (E)** Distribution of absolute tolerance indices for multi-digit modulating electrodes for each participant. **(F)** Bar plots summarizing the number of modulating electrodes for each digit (n=140). **(G)** Bar plots summarizing the number of modulating electrodes exhibiting excitatory or inhibitory responses (n=65) for each participant. **(H)** Arrays maps showing tuning of the channels across all digits for participants C1 and P5.

In the case of multi-digit RFs, single-unit analysis was performed in participant C1 to determine whether different neurons were selectively tuned to different digits or if the same neuron has multiple RFs spanning multiple digits. From the 64 electrodes in BA1, we isolated 10 individual units **(Supp. Fig. 3)**. Units were extracted based on distinct waveform shapes and their clustering in principal component space (**Supp. Fig. 3D**). For each unit, firing rates aligned to touch onset were computed, similarly to the multi-unit analysis (**Supp. Fig. 3A**). Among these units, 5 showed significant modulation in response to tactile stimulation (**Supp. Fig. 3B**) and consistently with the multi-unit activity measured in C1, most of the responsive single units showed multi-digit activation (**Supp. Fig. 3C**).

### Receptive fields of BA1 span the digits from tip to base

To characterize how the tactile information is processed within BA1 under the specific tactile stimulation applied, we analyzed both spatial and temporal neural responses evoked by actual touch with visual feedback in participant C1 and P5. For each modulating electrode and digit, we extracted the peak response time, and we predicted the RF centroids (predicted RFs) along the finger (**Fig. 3A, Supp. Fig. 4A**). We found that as the RF location shifts from the fingertip toward the base of the finger, the peak firing rate occurs progressively later. In C1, most RF centroids were located near the fingertip, while in P5 RF centroids were more spread across the finger. Overall, the spatial organization of responses reveals a systematic progression of peak firing times across electrodes along the digit. This analysis also allowed us to visualize how evoked activity propagates across this cortical region of SC and how the individual fingers (D2 and D3) are oriented within the targeted area (**Fig. 3B, Supp. Fig. 4B**). To validate the predicted RF centroids in C1, we then compared them with the centroids of the projected fields (PFs), defined as the patch of skin perceived by the participant when electrically-stimulating with the same electrode^27^. This comparison revealed a significant correlation between the two measures (C1: D2: 0.60; D3: 0.64, *p<0*.*05*), suggesting that the spatial distribution of the tactile-evoked neural activity is similar to the one elicited via intracortical stimulation (ICMS) (**Fig. 3C**).

**Fig. 3.**
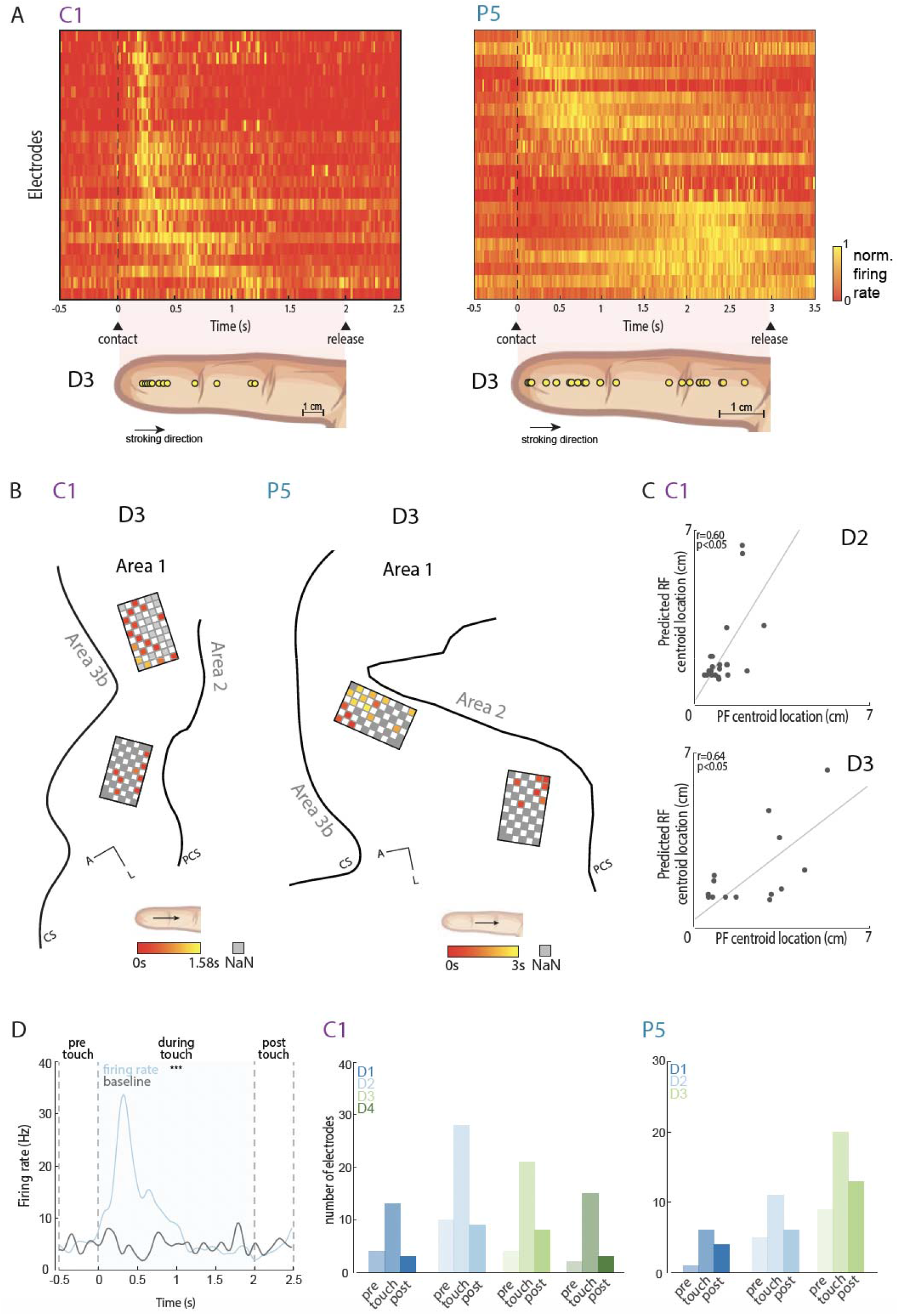
Spatiotemporal dynamics in response to touch. **(A)** Temporal progression of peak neural responses across modulating electrodes for digit D3 in participants C1 and P5. Electrodes are ordered according to their peak response time, and the predicted RF centroids along the digit are shown. Yellow indicates the time of the peak firing rate. **(B)** Propagation of touch-evoked activity across the cortical region of the SC for digit D3 in participants C1 and P5. Electrode colors represent the peak response time, with earlier responses shown in red and later responses in yellow. **(C)** Relationship between predicted RF centroids and PF centroids for participant C1 for digits D2 and D3. Each point represents one electrode. **(D)** Left: example of smoothed firing rate from a sensory array electrode in response to a stimulation of a single digit (light blue) and baseline firing rate (black line). Three-time windows of interest are highlighted: pre-touch, during touch and post-touch phases. Shaded areas represent firing rates significantly different from baseline. Right: summary of the number of electrodes showing significant neural responses compared to baseline for each digit and time window for participant C1 and P5.

### Neural responses in BA1 are time-locked to tactile events

We next examined the dynamics of the responses over time using a rate-coding analysis. Firing rates were computed across three-time windows (pre-touch, during-touch and post-touch phases) and compared with baseline activity (no-touch). Results revealed robust neural responses that were time-locked to tactile events, with most electrodes showing firing rates significantly different from baseline only during the touch phase (In C1, D1: n=12, D2: n=28, D3: n=21, D4: n=15; in P5, D1: n=6, D2: n=11, D3: n=20, during touch phase, *p<0*.*05*) **(Fig. 3D)**. Neural activity increased following touch onset and returned to baseline levels during the pre-touch and post-touch phases. Additionally, we assessed the consistency of the responses across multiple sequentially delivered tactile events (n=3). Population activity showed comparable response across the three consecutive touches (**Supp. Fig. 5A**). Indeed, neural activity did not show significant adaptation across repetitions (C1: *p=0*.*68;* P5: *p=0*.*62;* Friedman’s ANOVA; **Supp. Fig. 5B**), indicating consistent responses across touches.

### Motor cortex encodes tactile responses in absence of movement

While responses to tactile stimuli are well established in SC, they remain understudied in MC. Consequently, we analyzed neural responses in the MC elicited by physical touch with vision. Despite the absence of active movement, a substantial proportion of cortical neurons exhibited significant modulation during the task (see Methods - Multi-unit analysis). We calculated the normalized firing rates for each electrode responding to mechanical stroking of the four digits in C1 and C2 and three digits in P5 (**Fig. 4A**). Focusing on the most active array (the lateral array), 84% of the electrodes exhibited significant modulation in C1, 58% in C2, and 23% in P5 (**Fig. 4B**).

**Fig. 4.**
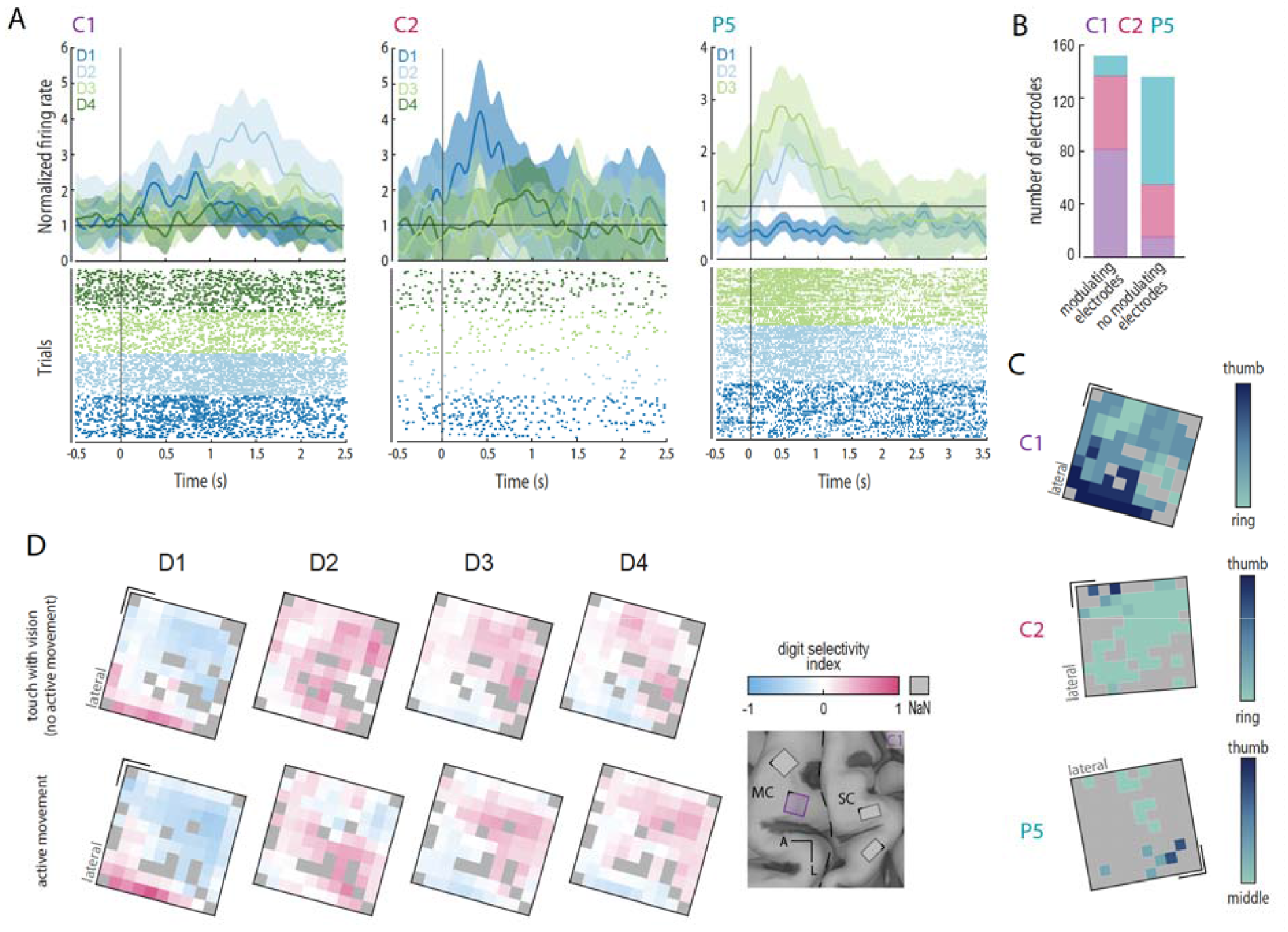
Touch-evoked responses in the human motor cortex (MC). **(A)** Example of smoothed normalized firing rates and raster plots from motor array channels in each participant in response to stimulation of each digit. Shaded areas indicate standard deviation. Black vertical line indicates touch onset. **(B)** Bar plot summarizing the total number of modulating electrodes from the motor arrays for each participant (n=288). **(C)** Digit selectivity gradient maps for each participant. Dark blue indicates channels responding more strongly to tactile stimulation of the thumb and index finger, while light blue indicates electrodes responding more strongly to stimulation of the middle and ring fingers. **(D)** Maps of digit selectivity index for each electrode of the motor arrays during tactile stimulation of different digits (top) and during attempted individual digits movements (bottom) in C1. Pink indicates higher neural activity for a given digit at a given electrode compared with the other digits recorded from the same electrode. Non-modulating electrodes are shown in grey.

To characterize the spatial organization of MC responses, we calculated the digit selectivity gradient for each electrode by computing the difference between the activation evoked by the touch of a digit and the mean activation across all four digits, normalized by the mean activation across digits^16^. For each electrode and digit, we then computed the Spearman correlation between the digit selectivity value and the digit identity. This analysis revealed a spatially and somatotopically organized pattern of activation (**Fig. 4C-D; Supp. Fig. 6**), even in the absence of active movement. This means, for example, that mechanical touch applied to the thumb elicited activity in neurons corresponding to the thumb representation in the MC.

To quantify the interplay between MC and SC, we compared the activity of MC during active movement and passive touch. Specifically, the tactile-evoked maps were compared with those obtained during individual digit movements, in which participants were asked to attempt to make finger movements following audiovisual instructions (for further details, see Emonds et al.^29^). This approach allowed a direct comparison of digit-specific activations across conditions. We found a significant similarity between the spatial pattern of MC activations in the active and passive conditions (Pearson’s correlation: C1: D1=0.92, D3=0.78, D4=0.66, *p<0*.*001*; C2: D1=0.31, D2=0.48, D3=0.36, D4=0.45, *p<0*.*05*; P5: D3=0.89, *p<0*.*05*), indicating that touch with visual input activates digit-specific representations in MC that closely resemble those activated during attempted movement in all participants (**Fig. 4D and Supp. Fig. 6**).

### Vision does not modulate somatosensory and motor responses evoked by touch

To determine whether visual input might modulate the tactile-evoked activity in either SC or MC, we recorded cortical activity during touch delivered without visual feedback (hidden touch). Participants were blindfolded and therefore unable to predict when touch would occur. Neural activity in this condition was directly compared with the activity recorded during touch with vision. We first calculated the normalized firing rates from all the electrodes in SC (**Fig. 5A**), during both touch with and without vision, for each participant. In SC, we analyzed the average neural responses of electrodes tuned to both conditions. Indeed, the average peak firing rates of modulating electrodes were highly correlated between the two conditions (Pearson’s correlation: C1: r=0.78, *p<0*.*001*, C2: r=0.98, *p<0*.*001*; P5: r=0.92, *p<0*.*001*; **Fig. 5B**). Within each digit, peak firing rates were comparable across conditions, indicating comparable response magnitudes regardless of visual inputs.

**Fig. 5.**
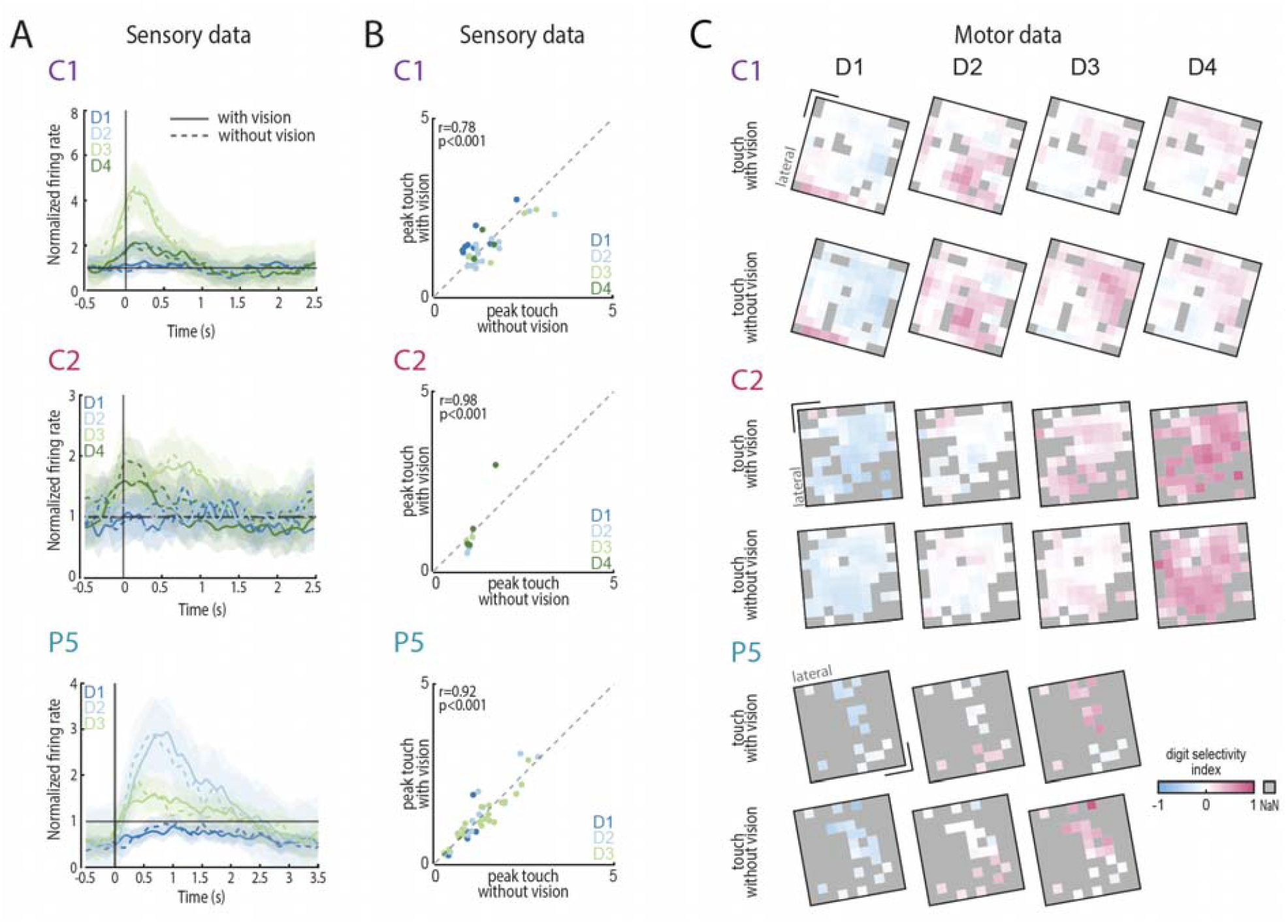
Touch-evoked response in absence of vision. **(A)** Example of smoothed normalized firing rates from sensory array channels in each participant in response to stimulation of each digit. The solid line represents the touch with vision condition, and the dotted line represents the touch without vision condition. Shaded areas indicate standard deviation. Black vertical line indicates touch onset. **(B)** Relationship between peak firing rates during the touch with vision and without vision conditions for each participant. Each dot represents one electrode modulating for a specific digit. **(C)** Maps of digit selectivity index for each electrode of the motor arrays during tactile stimulation of different digits with vision (top) and without vision (bottom) for the three participants. Pink indicates higher neural activity for a given digit at a given electrode compared with the other digits recorded from the same electrode. Non-modulating electrodes are shown in grey.

In absence of vision, evoked population responses were also observed in MC. To investigate MC activation during corporeal touch with and without vision, we compared the modulation maps obtained in each condition. For each digit, the spatial activation of the cortical maps showed high correlation between vision and no-vision conditions (Pearson’s correlation: C1: D1=0.95, D2=0.93, D3=0.92, D4=0.71, *p<0*.*001*; C2: D1=0.63, D2=0.62, D3=0.43, D4=0.62, *p<0*.*01*; P5: D1=0.62, D2=0.86, D3=0.80, *p<0*.*05*; **Fig. 5C**). Additionally, given the gradient of digit preference for the two conditions (**Supp. Fig. 7A**), the similarity between the maps remained significantly higher than chance (Pearson’s correlation C1: r=0.89, *p<0*.*001*, C2: r=0.92, *p<0*.*001*, P5: r=0.89, *p<0*.*001*). Significance was further assessed using a permutation test in which digit labels were randomly shuffled 10,000 times to generate a null distribution of correlation values (**Supp. Fig. 7B**). This indicates that MC exhibits a spatially organized and somatotopic pattern of activation independently from the visual inputs.

### Observing someone else being touched does not elicit significant neural response in SC and MC

To investigate whether SC and MC exhibit vicarious activity, we examined the observed-touch condition in which touch was applied to another person. The total number of modulating electrodes in both SC and MC was substantially smaller than during touch delivered on the participants’ hand. For participant C1, only 1% of the electrodes showed modulation in SC and 9% in MC. For participant C2, 9% of the SC electrodes and 47% of MC electrodes exhibited modulation, whereas for participant P5 11% of electrodes were modulated in SC and 6% in MC (**Fig. 6A**). Participant C2 exhibited stronger evoked activity in the MC, with response patterns that resembled those observed during actual touch (**Fig. 6B, middle**), compared to C1 and P5.

**Fig. 6.**
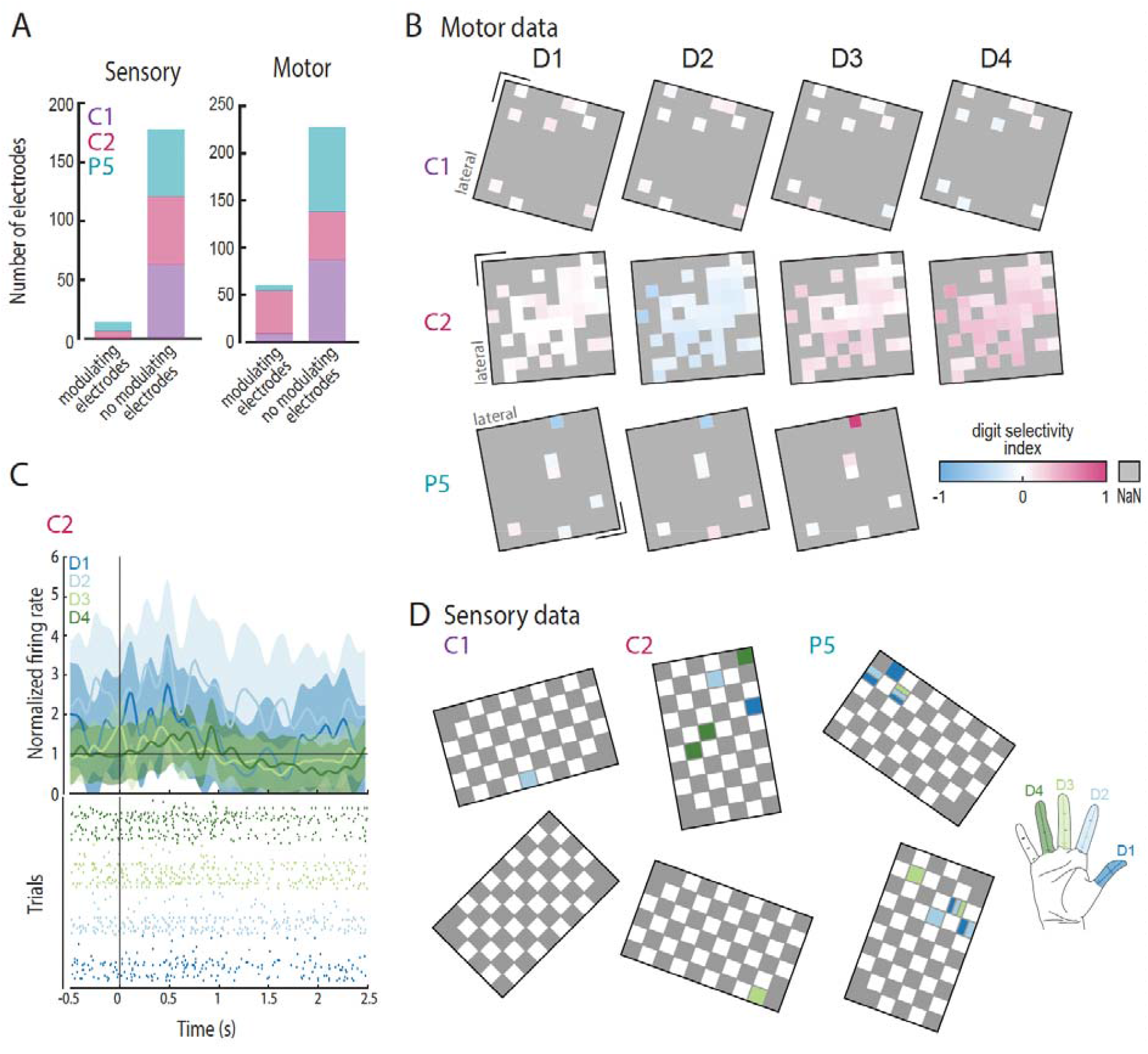
Vicarious activity in the human sensorimotor cortex (SC and MC). **(A)** Bar plots summarizing the total number of modulating electrodes in the SC (n=192) (left) and MC (n=288) (right) in the three participants. **(B)** Maps of digit selectivity index for each electrode of the motor arrays during observation of touch of different digits for the three participants. Pink indicates higher neural activity for a given digit at a given electrode compared with the other digits recorded from the same electrode. Non-modulating electrodes are shown in grey. **(C)** Example of smoothed normalized firing rates and raster plots from a sensory array channel in participant C2 in response to observation of touch of each digit. **(D)** Arrays maps showing tuning of the channels across all digits for the three participants.

Considering the modulating electrodes in this condition, we computed the firing rates and found weak evoked responses that lacked event-locked modulation to touch onset (**Fig. 6C**). To further support that observed touch does not elicit robust neural responses, we focused on modulating electrodes in touch with vision condition and extracted the maximum normalized firing rate during both touch with vision and observed touch. We then compared the resulting distributions separately for the sensory and motor arrays. This analysis further supports the overall absence of significant responses during observed touch, as the distributions show reduced activity, with maximum firing rates during observed touch being around baseline levels **(Supp. Fig. 8)**. Statistically significant differences between conditions (Wilcoxon signed rank test, *p<0*.*05*) were found for all participants **(Supp. Fig. 8)** indicating weaker responses during observed touch compared to touch with vision. Additionally, given the overall small proportion of modulating electrodes during the observation of touch on another person’s hand, no evidence of somatotopic was found in SC for all participants and in MC for C1 and P5 (**Fig. 6B-D**).

### Touch location can be decoded from the neural activity of both SC and MC

To investigate the neural response at network level, we calculated neural manifolds in both SC and MC^30^. We performed PCA on digit-averaged z-scored firing rates of all electrodes (**Fig. 7A-B, Supp. Fig. 9**). For participants C1 and P5, trajectories in the PC space corresponding to individual fingers were well separated both when considering MC and SC activity during touch with and without vision. In contrast, in the observed touch condition, trajectories corresponding to different digits largely overlapped.

**Fig. 7.**
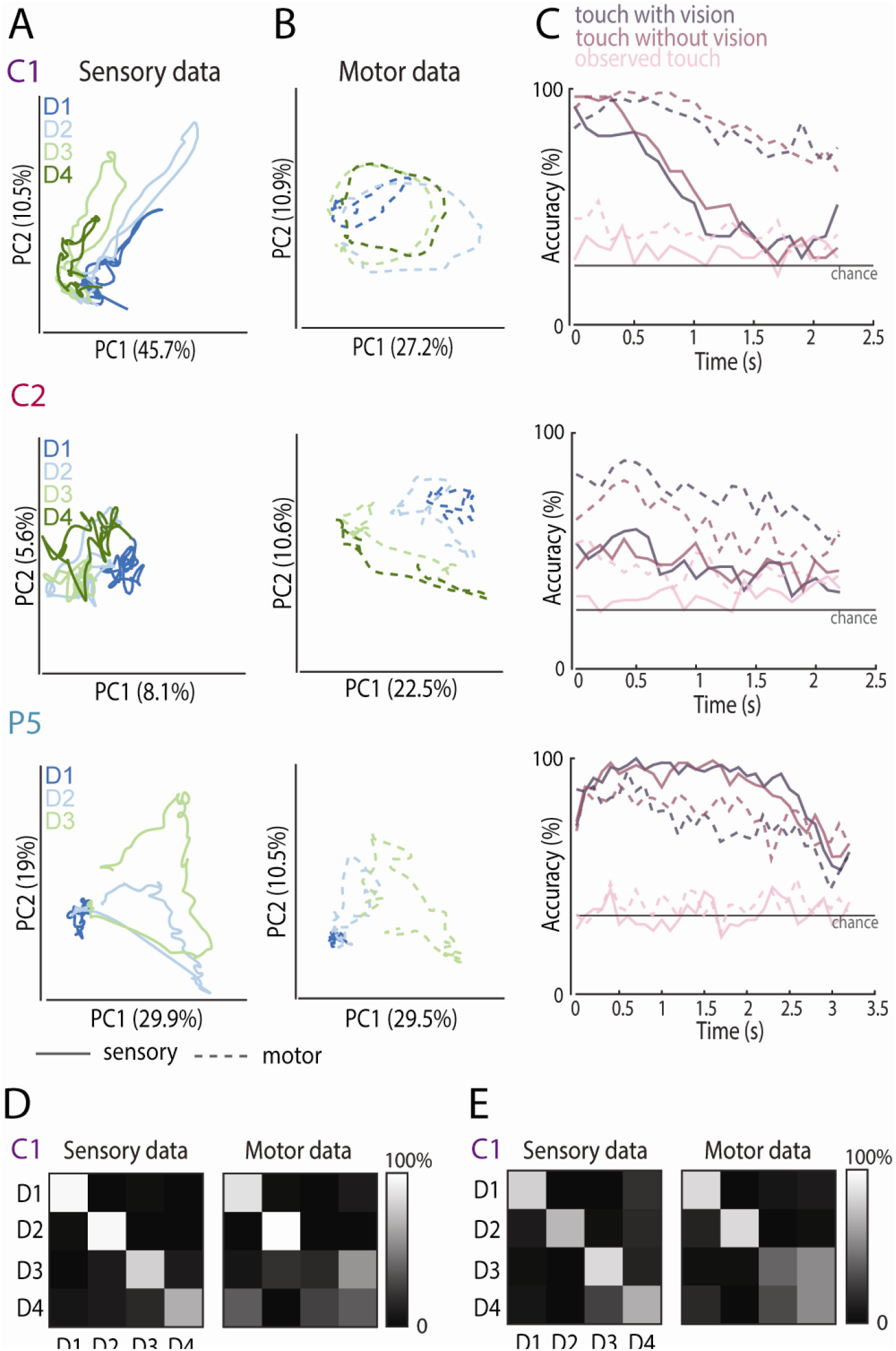
Cortical digit representation during touch. **(A)** Neural activity from the sensory arrays during tactile stimulation with vision projected into a 2D PC space for the three participants. Each color represents one digit. **(B)** Neural activity from the motor arrays during tactile stimulation with projected into a 2D PC space for the three participants. Each color represents one digit. **(C)** Decoding performance over time for each condition, computed separately for the sensory and motor arrays. The dotted line represents decoding accuracy using motor data and the solid line represents decoding accuracy using sensory arrays data. The horizontal line indicates chance level. Each condition is color-coded. **(D)** Confusion matrices obtained using a model trained on the touch with vision condition and tested on the touch without vision condition for C1 during touch window (D1-D4 for C1). **(E)** Confusion matrices obtained using a model trained on the touch without vision condition and tested on the touch with vision condition for C1 during touch window (D1-D4 for C1).

In addition, we classified which digit has been touched within each condition and across modalities using z-scored firing rates computed during the touch time window. For both SC and MC arrays, the classification performance was higher than chance during touch with and without vision. In participant C1, classification accuracy was 54.1±20.1% in SC and 83.8±9.4% in MC during touch with vision, and 60.6±24.3% in SC and 86.4±11.5% in MC during touch without vision; in participant C2 it was 48.8±10.9% in SC and 65.8±14% in MC during touch with vision, and 44.6±6% in SC and 62.2 ±10.8% in MC during touch without vision; and in participant P5, it was 87.3±14.3% in SC and 72.4±11.7% in MC during touch with vision, and 86.5±12.4% in SC and 76.1±7.3% in MC during touch without vision (**Fig. 7C**). In contrast, decoding accuracy in the observed condition was almost at chance for all three participants, particularly when considering neural activity recorded from SC (**Fig. 7C**).

Given the similarity of neural responses between touch with and without visual feedback, we additionally performed cross-condition decoding using the combined dataset. Specifically, an SVM classifier was trained on one condition and tested on the other. Decoding results showed that digits could still be correctly classified across conditions, indicating consistent neural representations across the two modalities. In participant C1 decoding accuracy was 75±7.9% in SC and 65±21.9% in MC for vision when trained on no vision; 85±14.5% in SC and 60±39.8% in MC for no vision when trained on vision. In participant C2 decoding accuracy was 35.8±15.2% in SC and 40.8±18% in MC for vision when trained on no-vision, while 32.5±11.4% in SC 40.8±23.6% in MC for vice versa. In P5 decoding accuracy was 100% in SC and 93.5±1.6% in MC for vision when trained on no-vision, while 100% in SC and 97.2±2.8% in MC for no vision when trained on vision (**Fig. 7D-E**; **Supp. Fig. 10A**). This similarity was also reflected in the PC space computed from the combined dataset, where digit trajectories from the two conditions followed overlapping and similar neural trajectories (**Supp. Fig. 10B**). These findings indicate that neural activity evoked during passive touch is largely invariant to visual feedback.

## Discussion

### Neurons in BA1 show multi-digit response, locked to touch-events

In this study, we investigated the population activity of human cortical neurons in SC during touch with visual feedback and its tuning properties. Among the modulating intracortical electrodes, approximately half of them responded to mechanical stimulation of more than one digit, reflecting in a high number of multi-digit RFs in BA1. This result is consistent with previous studies in NHPs showing that, compared to BA3b, neurons in BA1 more frequently exhibit multi-finger RFs, whereas such responses are only rarely observed in area 3b^8,9,31^. Interestingly, multi-digit electrodes maintained similar firing rates across digits and did not show strong digit selectivity, suggesting that in the case of multi-digit responses, the neuronal activity in BA1 is tolerant rather than highly selective, preserving similar levels of activity across different digits. Looking at the single-neuron level, the significant modulation in SC also shows a predominantly multi-digit response. Therefore, the RFs of these individual neurons span multiple digits, and within the somatosensory hierarchy they receive and integrate inputs from lower-level neurons with single-digit RFs.

Mapping the tactile RFs across the arrays revealed the expected somatotopic organization of BA1^32,33^, for all participants. Participant C2 presented more severe sensory impairments, which was reflected in a lower number of modulating electrodes in SC, and the less evident somatotopic organization; D4 was the digit with the lowest tactile perception threshold (**Supp. Table 1**) and showed the strongest evoked neural activity. In all participants, adjacent digits were represented in continuous cortical regions, with a progression from lateral to medial corresponding to thumb to little finger. This spatial organization confirms that BA1 preserves the topographic mapping of the digits^7,11^. While similar somatotopic organization has been reported mostly in NHPs^34–36^ or brain imaging studies^37–39^, and is consistent with human ICMS studies^40^, the present results extend these findings by directly characterizing neural responses to mechanical stimuli at a much finer spatial resolution, enabled by intracortical recordings with hundreds of microelectrodes. In addition, the separability between digits was further supported by the network analysis, which showed that individual digits could be reliably distinguished from SC neural activity. Although some electrodes exhibited multi-digit responses with similar firing rates across digits, population-level decoding was still able to identify the stimulated digit. This suggests that digit-specific information is distributed across the neural population, with neurons showing more selective responses contributing to determining which digit was touched, even when individual recording electrodes exhibit multi-digits RFs. Moreover, most RFs were located on the fingers, and indeed the arrays were positioned toward BA2 (the posterior edge of BA1) in all participants. This pattern is consistent with the organization described in NHPs^41^. However, it is difficult to clearly determine the orientation of the fingers within BA1. To assess individual digit representation, we quantified the spatial propagation of responses across the skin and the arrays during stroking. In NHPs, distal finger segments are typically represented near the border with BA2, whereas the finger bases map closer to area 3b (the anterior edge of BA1). In our participants, however, we did not observe this clear organization, due to the difficulty in precisely identifying areal boundaries.

Looking at the evoked neural dynamics in BA1, most of the modulation emerged following touch onset, whereas neural activity during the pre-touch and post-touch periods tended to return to baseline levels. These results indicate that SC responses are strongly time-locked to tactile events, consistent with previous reports^23^, excluding the presence of sensory expectation signals or neural adaptation to consecutive touches. For the latter, the timescale of the mechanical stimulation may be a contributing factor.

### Passive touch evoked somatotopic neural response in MC even in absence of movement

During touch with visual feedback, we observed widespread neural activity in the MC, even if the task did not involve any active movement. Importantly, analysis of the spatial activation patterns revealed a spatial and somatotopic representation of the digits in MC similar to the organization previously reported during active movement, in which participants were asked to perform individual finger movements^29^. The presence of somatotopically organized responses in MC during passive tactile stimulation suggests a tight functional coupling between SC and MC. Notably, the evoked activity observed in SC during attempted individual finger movements was weaker and did not exhibit vivid somatotopic organization (**Supp. Fig. 11**), suggesting that the functional communication between these two cortices may be stronger from SC to MC than in the opposite direction.

The modulation of MC activity by the identity of the stimulated digit supports that tactile inputs retain spatial specificity as they propagate through sensorimotor circuits, in contrast to a less specific movement somatotopy previously reported in MC^42,43^. In support to our finding, a previous study showed that direct microstimulation of BA1 evoke somatotopically-organized neural activity in MC^16^. In addition, BA1 integrates inputs across digits and is well positioned to relay processed tactile information to motor regions. BA1 sends direct projections to BA4 and BA6d, as demonstrated by tract-tracing studies in NHPs^44^. These connections provide a structural substrate for the transfer of processed tactile information from somatosensory to motor circuits. The observed digit-specific responses in MC therefore likely reflect cortico-cortical interactions between SC and MC that preserve somatotopic organization. In parallel, thalamo-cortical pathways may provide an additional route for sensory signals to influence MC^10^. Together, these results support the view that MC encodes sensory-related information during natural touch, even in the absence of movement. In an active context, sensory feedback from a given digit may preferentially inform the motor control of the same digit, supporting sensorimotor coordination^45^. The neural basis of sensorimotor interaction has been mostly studied in animals^46^, but still their role during exact tactile coding remains less explored. The present findings provide insight that digit-specific tactile information can be represented in MC even in the absence of movement, highlighting the coupling between the two cortices even during passive touch.

### Visual input does not modulate touch-evoked activity in both SC and MC

To investigate the contribution of visual information, we examined neural responses during touch both with and without visual feedback. In both conditions, we observed strong and well-defined neural activity in both cortices across the different digits. The magnitude of the overall spatial response was comparable across the three participants in the two conditions. This result indicates that the vision of the tactile stimulation does not modulate significantly the responses of cortical neurons in SC and MC, suggesting that visual input may not be strongly integrated at this stage of somatosensory processing. Indeed, the activity recorded in BA1 may primarily reflect the bottom-up sensory processing driven by the peripheral somatosensory inputs. This interpretation supports that early somatosensory areas are primarily driven by direct tactile inputs, whereas stronger visual-tactile integration typically emerges in higher-order multisensory regions (secondary SC)^47^. This is further validated by the finding that classifiers trained on one condition (touch with vision or touch without vision) and tested on the other achieve high accuracy in distinguishing stimulus locations. Therefore, digit information is encoded in SC and MC regardless of the presence of visual input. Our finding are in contrast with a previous study^23^ recording from SC, which reported weaker neural responses when touch was delivered without visual feedback, suggesting that visual inputs might modulate SC activity, making it stronger. Several factors may account for this discrepancy, including inter-subject variability, variability in the level and extent of SCI, and differences in the specific cortical regions targeted by the microelectrode arrays. In that study, tactile stimulation was delivered to the arm at a location where the participant reported numbness, which may have influenced the observed neural responses.

Moreover, the hidden touch condition further allowed us to assess whether the activation measured in MC was driven by visual cues related to the experimenter’s hand movement approaching the participant’s digits, potentially engaging mirror-like responses, or by the tactile sensory inputs generated by the stimulation. The presence of similar responses in both conditions argues against purely visually driven explanation and instead suggests that MC activity is related to the sensory consequences of the touch, by receiving sensory-related inputs. However, whether these inputs reach MC through thalamocortical pathways or via corticocortical projections remains to be clarified.

### BA1 neurons do not show vicarious activity

Previous literature of touch observation has primarily relied on brain imaging techniques, which provide limited spatial resolution and therefore make it difficult to precisely localize the cortical regions in which there is activity during observed touch. In contrast, the results presented here provide high-spatial-resolution recordings from specific regions of MC and SC, allowing us to more directly assess the impact of visual inputs on neuron populations within these cortical areas. During the observation of a tactile stimulus applied to another person’s hand, very few electrodes showed significant response in SC or MC. The evoked neural activity of the few modulating electrodes did not exhibit a defined spatial organization or event-locked modulation in either SC and MC, suggesting that the vicarious activity is weak and not differentiated across fingers. Consistent with this finding, the different digits were not represented in a spatially-unique neural space. Nevertheless, the discriminability between fingers was higher in MC than in SC. One possible explanation for this effect is the involvement of the mirror neuron system, which may generate motor-related representations during the observation of actions performed by others^48,49^. Indeed, the array’s placement in BA4 and BA6d includes networks classically reported as part of the mirror neuron system.

Overall, these results suggest, in line with previous work^23^, that observation of touch applied to another individual is not sufficient to elicit strong neural responses in SC or MC. This finding further supports the idea that visual information related to observed touch may be integrated at higher stages of the somatosensory processing hierarchy, as proposed by previous studies^50,51^. Evidence from studies of higher stages of tactile processing, in particular on the posterior parietal cortex (PPC), indicates that neurons in this region respond to inputs from multiple sensory modalities, particularly visual signals^51^. This suggests that PPC may play a key role in multisensory integration and sensorimotor planning^4^, and therefore could be engaged during the observation of touch^50,51^.

### Implications for brain computer interfacing

The use of intracortical microelectrode arrays implanted in sensorimotor cortical areas offers a unique opportunity not only to restore tactile sensations through ICMS, but also to deepen our understanding of the neural mechanisms underlying touch perception. Investigating how tactile information is encoded in the SC, including its spatial organization, temporal dynamics, and interactions with visual inputs, can provide useful insights for the design of BCIs aimed at delivering artificial sensory feedback^52^. Understanding how tactile signals are represented within the SC, and specifically within the hand area, is a crucial step toward developing biomimetic stimulation strategies^53,54^. Such strategies aim to recreate patterns of neural activity that closely resemble those occurring during natural touch, thereby improving the quality and usability of sensory feedback delivered through ICMS^55,56^. Characterizing the spatiotemporal properties of tactile responses in this area therefore provides important information for the development of encoding algorithms designed to evoke more naturalistic percepts.

In the present study, we also investigated neural responses to observed touch that could in principle provide useful information for guiding microelectrode array placement. This question is particularly relevant in clinical contexts involving deafferented patients, where the absence of peripheral sensory input makes it more challenging to localize body-part representations in the SC. Activating this region through observed touch has been considered a potential alternative. However, we did not observe significant activation in response to observed touch in the recorded areas. This suggests that visual observation of tactile events alone may not provide a reliable functional marker for targeting implant locations within BA1. Consequently, alternative strategies, such as relying on residual sensory responses or neural activity during attempted movement tasks^57^, may be required to guide implantation in such patient populations. Notably, this absence of modulation during the observation of touch may also represent a practical advantage for the design of ICMS strategies, as it suggests that artificially evoked tactile percepts are unlikely to be influenced by observing touch on others.

Finally, studying the functional interactions between the SC and MC has important implications for BCI development, especially in closed-loop scenarios^58^. A better understanding of how sensory and motor signals interact may contribute to improving the decoding of motor intentions and to the development of bidirectional neuroprosthetic systems in which motor commands and sensory feedback are continuously integrated.

### Limitations

The present study has some limitations that should be considered. First, the number of participants included in the study was limited (n=3), which is a common constraint in intracortical recording studies involving human participants implanted with microelectrode arrays. Second, tactile stimulation was delivered manually. This approach may introduce variability across trials in the force applied, contact characteristics, and timing of the stimuli. However, we did not observe differences in neural firing rate modulation as a function of the force applied during the stimulation (**Supp. Fig. 5C**), suggesting that variability associated with manual delivery did not substantially affect the results. Another limitation concerns the spatial coverage of the recordings. Neural activity was recorded from a relatively small area of the SC, specifically within the finger representations of BA1, as well as from a limited region of the MC. As a result, the present findings may not fully capture the broader organization of tactile representations across other regions.

## Methods

### Participants

This study was conducted under an Investigational Device Exemption from the U.S. Food and Drug Administration and approved by Institutional Review Boards at the University of Chicago and the University of Pittsburgh. The clinical trial is registered at ClinicalTrials.gov (NCT01894802). Informed consent was obtained before any study procedures were collected from all participants. Participant C1 (male), 55-60 years old at the time of implant, presented with a C4-level ASIA D spinal cord injury (SCI) that occurred 35 years prior to implant. Participant C2 (male), 60-65 years old at the time of implant, presented with C4-level ASIA D SCI and right brachial plexus injury that occurred 4 years prior to implant. Participant P5 (female), 51 years old at the time of implant, presented with a C8-level ASIA A SCI that occurred 6 years prior to implant.

Participants retained residual tactile sensation in regions corresponding to the receptive fields recorded by the implanted electrodes^59^. We used monofilaments^60^ tests to measure tactile thresholds before implantation (**Supp. Table 1**).

### Cortical implants

Four Neuroport electrode arrays (Blackrock Neurotech, Salt Lake City, UT, USA) were implanted in the left hemisphere of each participant. Two of the arrays (one medial and one lateral array) were placed in the hand representation of Broadmann’s area 1 of the SC (targeting layer 4). These two arrays were 2.4 mm x 4 mm, with 60 electrodes arranged in a 6x10 grid. Each electrode shank was 1.5-mm long. The electrodes were wired in a checkerboard pattern, such that 32 electrodes per array could measure neural activity. ICMS could also be delivered by each of these electrodes. The other two arrays were placed in the arm and hand representation areas of the MC. These arrays were 4 mm x 4 mm, with one-hundred 1.5-mm long electrode shanks wired such that 96 electrodes per array could measure neural activity. The inactive shanks were located at the corners of these arrays. Two percutaneous connectors, each connected to one sensory array and one motor array, were fixed to the participant’s skull. Array placement of the four arrays was targeted based on functional neuroimaging (fMRI or MEG) as the participants attempted to make movements of the hand and arm^57^, and imagining feeling sensations on their fingertips, within the constraints of anatomical features such as blood vessels and cortical topography (**Fig. 1A**).

### Experimental protocol

To investigate how touch and visual inputs are integrated in sensory processing, we focused on four finger locations, thumb (D1), index (D2), middle (D3), and ring (D4) for participant C1 and C2, and three finger locations (D1, D2, D3) for participant P5, on the contralateral hand across three experimental conditions. These digits were selected because the implanted electrode arrays were positioned over cortical regions associated with these digits, allowing us to record the neural activity related to tactile stimulation of these fingers. Three task conditions were administered to all participants: *“touch with visual feedback”*, in which the experimenter applied tactile stimulation to the participant’s fingers while the participant could see the scene; *“touch without visual feedback,”* in which the same tactile stimulation was delivered but the participant was blindfolded; and *“observed touch”*, in which the experimenter applied tactile stimulation to the fingers of their own left hand while the participant observed.

Across all conditions, the same type of tactile stimulus was done: a stroke applied from the tip to the base of the digit lasting approximately 2s for participant C1 and C2 and 3s for participant P5. Mechanical stimulation was delivered using either a cotton swab or a sensorized pen (ATI Nano17, USA). The sensorized pen enabled precise alignment of touch onset with neural recordings as well as the synchronized recording of the normal and tangential forces applied to the skin (6 components) during each stroke. When the cotton swab was used, a manual trigger was collected for each trial to allow alignment with touch onset. Each trial consisted of three consecutive strokes delivered to the same finger. In each session, ten trials were collected for each condition performed and each digit, resulting in a total of 30 trials for C1 and C2, and 36 trials for P5 per condition per digit. The order of digit stimulation was randomized. Throughout the experiment, participants were seated with their right hand comfortably placed, and the experimenter performing the tactile stimulation was positioned in front of them. Data collection was conducted in a manner that accommodated the time constraints and specific needs of participants.

### Neural recordings

Neural activity from SC and MC was recorded using the NeuroPort system (Blackrock Neurotech, Salt Lake City, UT, USA) at a sampling rate of 30 kHz. The neural signals were high-pass filtered with a first-order 750-Hz filter. Spiking events were detected when the signal crossed a threshold set at -4.5 time the root mean square (RMS) defined at the beginning of each recording session. Whenever this threshold was exceeded, a snippet of the waveform was stored.

For C1 data were collected across two recording sessions spanning 17 months. During the first session, touch with visual feedback and observed touch conditions were performed. During the second session, all three experimental conditions were recorded. In both sessions, data were collected in 5 blocks. For C2 data were collected in two sessions over 2 weeks. In the first session, the touch with visual feedback and the observed touch conditions were performed. In the second session, the touch with visual feedback and the touch without visual feedback conditions were recorded. As for C1, each session consisted of five data collection blocks. For P5 data were collected in one session, where all three conditions were performed.

### Multi-unit analysis

Multi-unit activity for each trial was aligned to the moment of contact (touch onset). In conditions using the sensorized pen, touch onset was determined from the pen’s force readings, whereas in conditions using the cotton swab, touch onset was extracted from the manual trigger. Firing rates for each electrode were computed using 20 ms bin size over a time window spanning from 0.5 s before touch onset to 2.5 s after touch onset (for P5 up to 3.5s after touch onset). The baseline firing rate was calculated for each trial using a time window from -3.5s to -0.5s before the first touch and then averaged across trials for each electrode, condition and finger. Firing rates for each finger were normalized within each electrode by dividing the firing rate by mean baseline activity of the corresponding digit and electrode.

Multi-unit responses were characterized using z-scored firing rate. For each electrode, firing rates were first z-scored on a trial-by-trial basis relative to baseline activity. The z-scored responses were then averaged across trials to obtain a mean z-score for each electrode, condition and digit.

An electrode was considered modulated to a specific condition and digit when the mean z-score exceeded the threshold (excitatory response) or was below the negative threshold (inhibitory response), where the threshold was defined as:

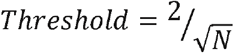

where N is the number of trials; for at least four consecutive time bins for participants C1 and P5, and three consecutive time bins for participant C2. This threshold accounts for the variability reduction introduced by averaging across trials and corresponds to two times the standard deviation criterion scaled by the number of trials. Additionally, for all touch conditions, modulating electrodes were further verified through visual inspection to maximize the accuracy of the electrode selection.

To verify the selectivity of multi-digit RFs across digits, we computed an absolute tolerance index. For each multi-digit electrode, its absolute tolerance^61^ was calculated as:

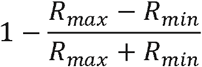

where *R*_*max*_ and *R*_*min*_ represent the maximum and minimum peak firing rates of the electrode across the modulated digits. The tolerance index ranges from 0 to 1, with a value of 1 indicating no variation in peak firing rate across digits.

### Single-unit analysis

Single sorting was performed offline using Plexon Offline Sorter (Plexon Inc., Dallas, TX, USA). Spike waveforms from each channel were imported into the software and visualized both in waveform shape and in principal component space. Units were identified on a channel-by-channel basis using the default sorting parameters provided by Plexon and manual inspection of waveform clustering. Putative single units were isolated based on the cluster separation in principal component space and waveform consistency. The quality of unit isolation was further assessed by examining inter-spike interval (ISI) distributions and cross-correlograms.

A subset of the analysis described above was repeated using the resulting isolated single units.

### Spatial-temporal dynamics for estimating RF centroids location

The spatial-temporal propagation of the neural activity across the electrode arrays was assessed by identifying, for each modulating electrode, the time at which the peak firing rate occurred during the stroking. The electrodes were sorted based on the peak times to visualize the signal propagation. For visualization purposes, we extracted the firing rate from 0.5 before touch onset to 2.5 s after touch onset for all modulating electrodes and we normalized each firing rate by its maximum value. Using the peak times, together with the average duration of the stroke and the length of each digit, we computed the predicted RF centroids. These predicted RF centroids were compared with the corresponding PF centroids for each electrode using Pearson correlation. In addition, to visualize the propagation and orientation of the finger within the array, electrodes were color-coded based on their peak timing.

### Task-selective activation

The temporal dynamics of the neural responses were performed through a rate coding analysis^62^. For each location, all modulating electrodes were considered. Three time-windows relative to touch onset were defined: pre-touch (-0.5 to 0 s), during touch (0 to 2 s), and post-touch (2 to 2.5s) for C1 and pre-touch (-0.5 to 0 s), during touch (0 to 3 s), and post-touch (3 to 3.5s) for P5. For each window, the mean firing rate was computed for each trial, finger and modulated channel. A mean baseline vector was also computed for each trial and time window. In each time window, statistical comparisons were performed between the mean firing rate and the baseline. Since the data was not normally distributed, Wilcoxon signed-rank tests were applied. The number of channels showing significant difference from baseline in each time windows was reported (*p<0*.*05*).

To assess whether adaptation occurred across the three touches, the data were divided into first, second, and third touch events. For all locations and modulated channels, firing rates were normalized by the baseline firing rate and averaged across trials. The maximum value of the normalized firing rate was then computed to obtain the peak firing rate. The distributions of peak firing rates were visualized, and a Friedman’s ANOVA test was performed to assess statistically significant differences between the peaks across the three touches, with the significance level set at *p < 0*.*05*.

### Digit selectivity gradient

We investigated the spatial pattern activation of the MC and SC by computing digit selectivity gradient maps. For each of the modulating electrodes, a digit selectivity index was calculated for each digit as the difference between the average firing rate for that digit and the mean firing rate across all digits, normalized by the mean firing rate across all digits. The resulting values were plotted across all channels for each digit to obtain digit-specific selectivity maps. Additionally, for each electrode and digit, we computed the Spearman correlation between the digit selectivity index and the digit identity (thumb=1, index=2, middle=3, ring=4). Electrodes exhibiting preferential responses to the middle and ring fingers yielded positive correlations, whereas electrodes preferring the thumb and index fingers yielded negative correlations.

To assess the similarity between conditions, we computed Pearson’s correlation between digit preferences profiles. Statistical significance was evaluated using a permutation test in which digit labels were randomly shuffled 10,000 times to generate a null distribution of correlation values.

### Digit decoding

For each condition (touch with vision, observed touch and touch without vision), neural activity was considered over the time window following touch onset. For each trial, firing rates were first z-scored relative to baseline. The firing rates were then divided into overlapping 300 ms windows with a 100 ms step size. Within each window, the mean z-scored firing rate across time was computed for each electrode and for each trial, resulting in a feature matrix of dimensions *trials x channels* for each time window. Digit identity was decoded using a support vector machine (SVM) classifier. For each time window, the feature matrix was split into training and test sets using 5-fold cross-validation, and classification accuracy was calculated by averaging performance across folds. This procedure was repeated across the entire window following touch onset. The resulting decoding accuracy over time was plotted for each condition to visualize the temporal evolution of digit-specific representations.

For the cross-decoding analysis, a SVM classifier was trained on neural activity from one condition (touch with or without vision) and tested on the other condition to predict the different digits. For each trial, the mean firing rate within the touch window was computed for each electrode and used as input features for the classifier. Classification performance was evaluated by computing the confusion matrix between the predicted and the true digit labels.

### Principle component analysis (PCA)

For each electrode, the average z-scored firing rate across trials was calculated using 20ms time bins from 0.5s before touch onset until 0.5s after the end of the mechanical stimulus. PCA was performed on the electrode-by-time matrix separately for the motor and sensory arrays, and for each experimental condition. Neural activity was then projected onto space defined by the first two principal components with the highest variance explained, to visualize the population dynamics during touch.

### Statistical analysis

All data were exported and processed offline in MATLAB (R2023a, The MathWorks, Inc., USA) and reported as mean value ± SD (unless elsewise indicated). All statistics were performed using the available built-in functions. Since the data did not meet the assumptions of normality, differences between firing rate distributions during tactile stimulation with the baseline (no stimuli) period were assessed using Wilcoxon signed-rank tests. Additionally, comparisons between repeated measurements across more than two groups (e.g., consecutive touches) were performed using the Friedman test, a non-parametric alternative for repeated measures. Correlations between variables (e.g., predicted RF centroids and PF centroids) were quantified using Pearson correlation coefficients unless otherwise stated. Additional details about the number of repetitions (n), and p-values for each experiment are reported in the results and in the corresponding figure legends.

## Acknowledgments

The authors are deeply grateful to the three subjects who freely donated months of their life for the advancement of knowledge and for a better future for people with SCI. We thank the whole CBRG team, especially Gary Blumenthal, for providing motor data for P5. We thank Prof. Sliman Bensmaia for his support during the initial phase of the study. The funder had no role in the experimental design, analysis, or manuscript preparation or submission. All authors had complete access to data. All authors authorized submission of the manuscript, but the final submission decision was made by the corresponding authors.

## Author contributions

M.C. performed all analyses, prepared the figures and wrote the paper; A.H.A. performed the experiments with C1 and C2; T.G.H. performed the experiments with P5; A.M.X.E. performed the individual digit movement experiments and performed the related analysis; A.S. supervised the motor data acquisition; R.A.G. supervised the clinical trial; C.M.G. programmed the experimental apparatus, supervised the project; G.V. designed the study, performed and supervised the experiments, supervised all the analyses, edited the figures and wrote the paper. All authors edited and proofread the manuscript.

## Data and materials availability

All data are stored at the Data Archive BRAIN Initiative. The code will be available on https://github.com/

