## Supplementary information for "Spatiotemporal encoding of touch signals in the human somatosensory and motor cortices"

**List of Supplementary Materials:**

Supp. Fig. 1. Somatotopic organization of digits within SC.

Supp. Fig. 2. Receptive field maps in SC.

Supp. Fig. 3. Single-unit responses in the human SC.

Supp. Fig. 4. Spatiotemporal dynamics in response to touch.

Supp. Fig. 5. Neural responses across consecutive touches

Supp. Fig. 6. Touch-evoked responses in the human MC.

Supp. Fig. 7. Touch-evoked response in absence of vision in MC.

Supp. Fig. 8. Maximum normalized firing rate distribution during touch with vision and observed touch

Supp. Fig. 9. Cortical digits representation during touch.

Supp. Fig. 10. Cross-decoding of digits between touch with and without vision.

Supp. Fig. 11. Attempted individual finger movements responses in the human SC.

Supp. Table S1. Tactile thresholds of the hand before implantation for each participant.

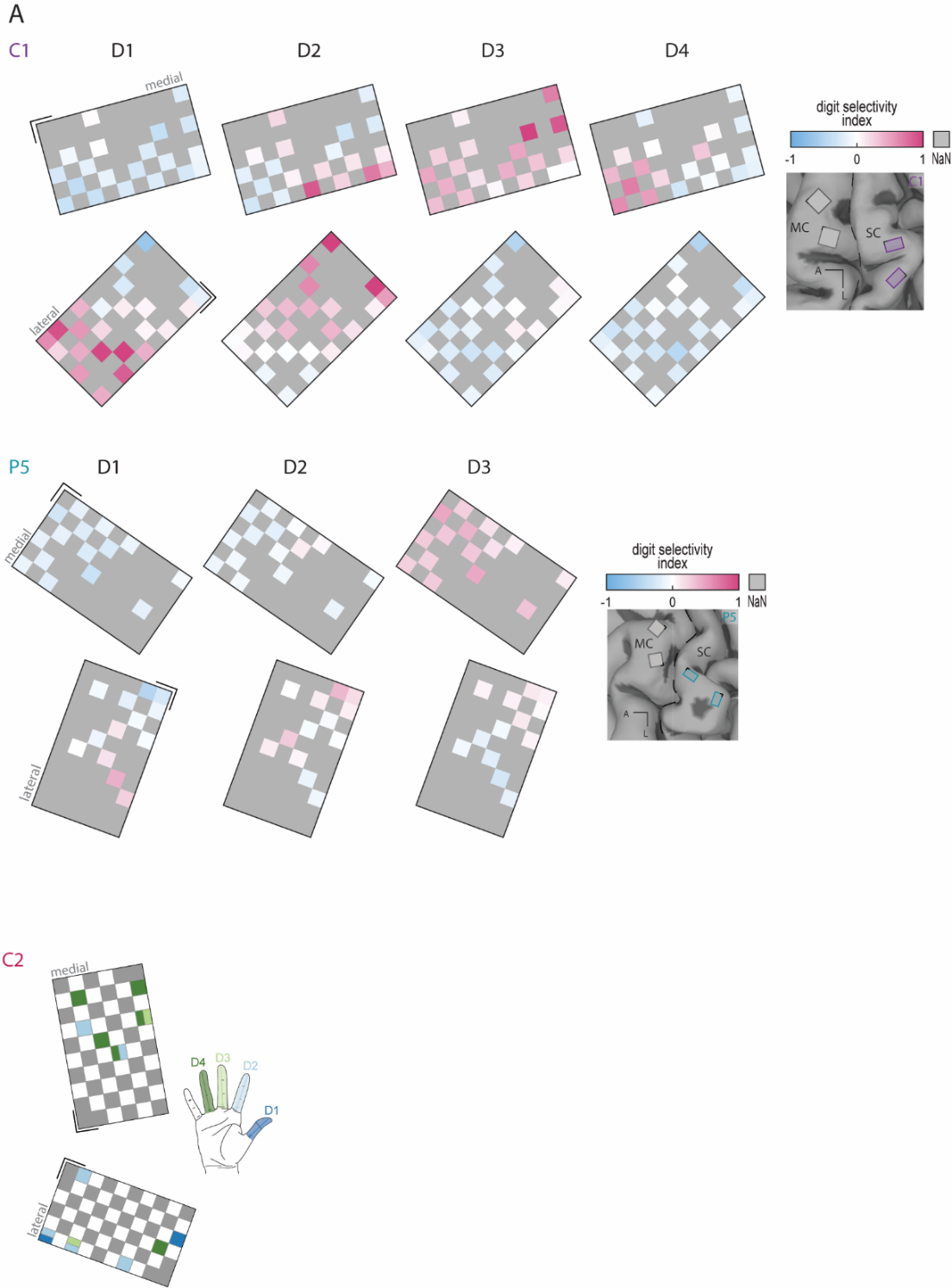

**Supp. Fig. 1. Somatotopic organization of digits within SC.** Maps of digit selectivity index for each electrode of the sensory arrays during tactile stimulation of different digits in C1 (top) and P5 (bottom). Pink indicates higher neural activity for a given digit at a given electrode compared with the other digits recorded from the same electrode. Non-modulating electrodes are shown in grey. MRI images for each participant with the related arrays’ placements are shown on the right. The arrays considered in this analysis are depicted in purple or green.

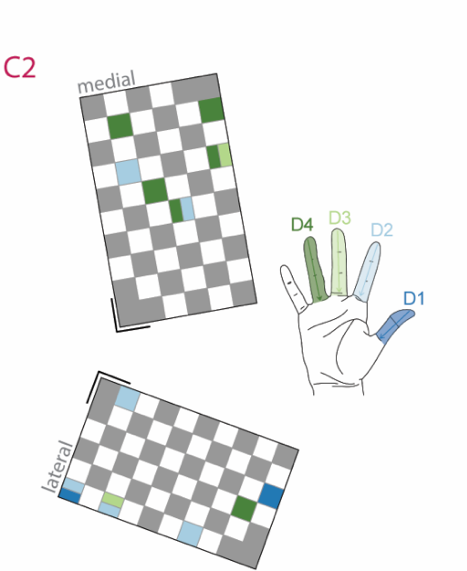

**Supp. Fig. 2. Receptive field maps in SC.** Map of the two sensory arrays showing the digit tuning of each electrode for participant C2.

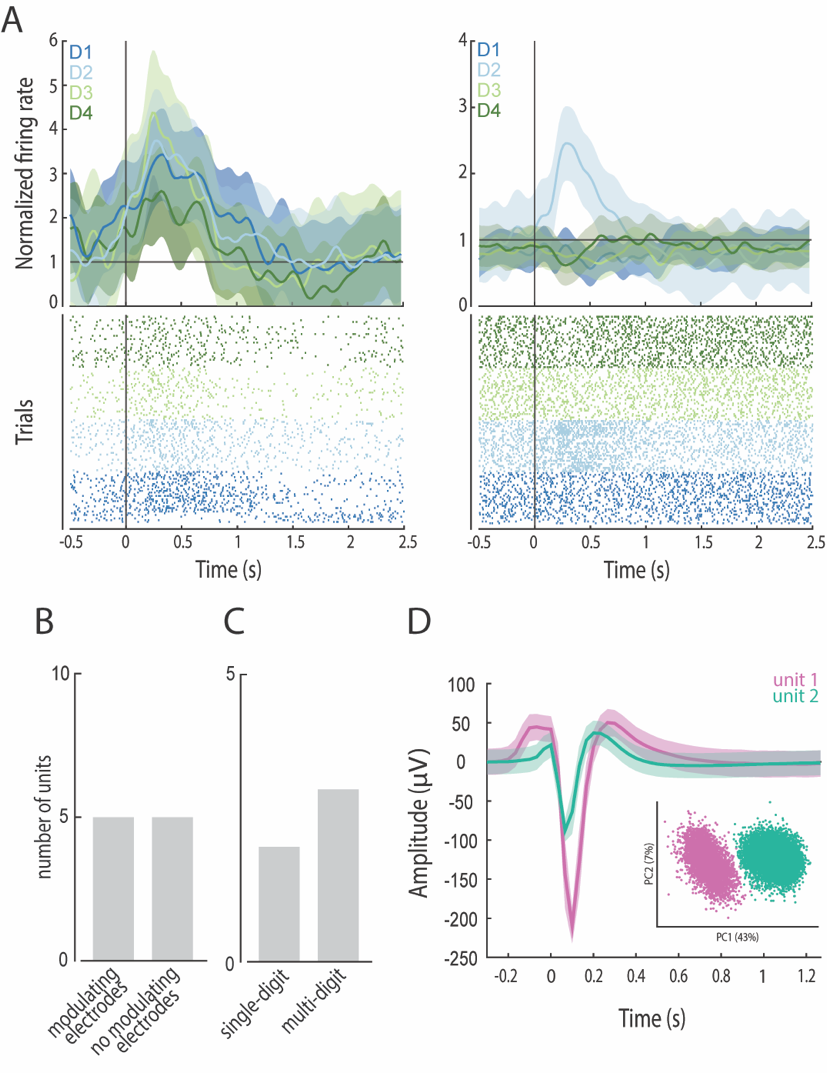

**Supp. Fig. 3. Single-unit responses in the human SC. (A)** Example of smoothed normalized firing rates and raster plots from two single units in response to stimulation of each digit. Shaded areas indicate standard deviation. Multi-digit activation (left) and single-digit activation (right) are displayed. Black vertical line indicates touch onset. **(B)** Bar plot summarizing the total number of modulating single units (n=10). **(C)** Bar plot summarizing the number of multi- and single-digit tuned single units (n=5)**. (D)** Example waveforms of two units extracted from a single electrode and their clustering in principal component space.

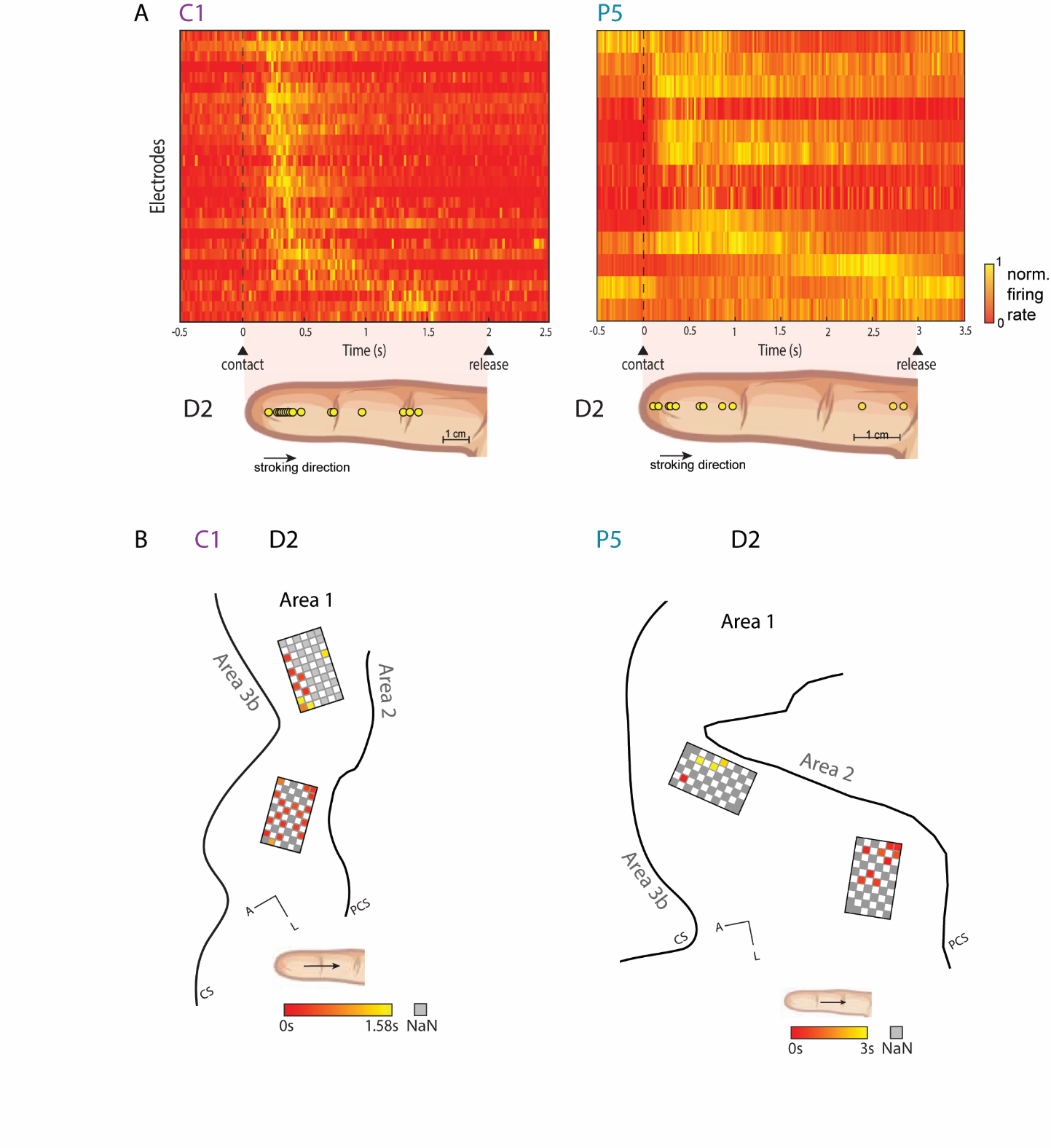

**Supp. Fig. 4. Spatiotemporal dynamics in response to touch. (A)** Temporal progression of peak neural responses across modulating electrodes for digit D2 in participants C1 and P5. Electrodes are ordered according to their peak response time, and the predicted RF centroids along the digit are shown. Yellow indicates the time of the peak firing rate. **(B)** Propagation of touch-evoked activity across the cortical region of the SC for digit D2 in participants C3 and P5. Electrode colors represent the peak response time, with earlier responses shown in red and later responses in yellow.

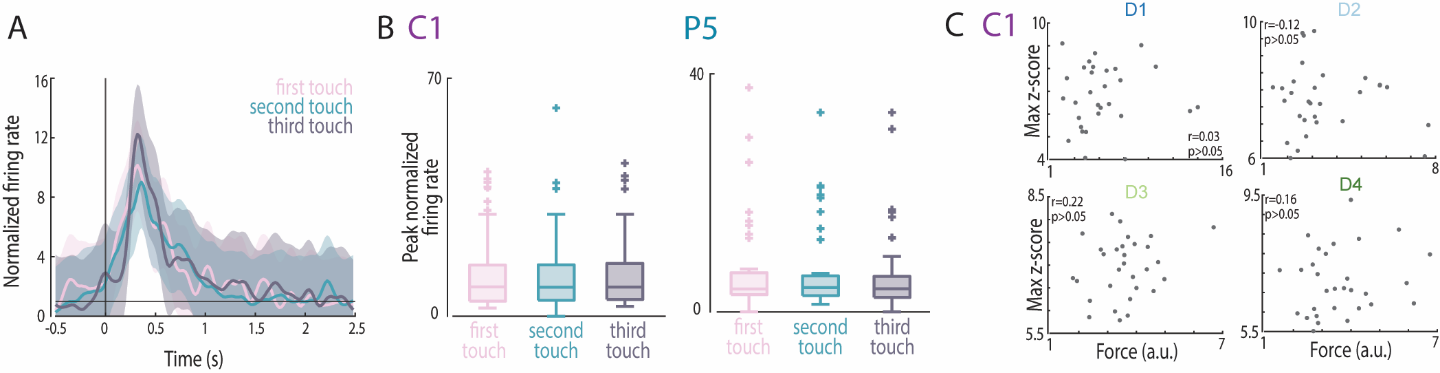

**Supp. Fig. 5. Neural responses across consecutive touches. (A)** Example of normalized firing rates across 3 consecutive touches from a single sensory array electrode. Black vertical line indicates touch onset. Each touch is color-coded (first touch = pink, second touch = turquoise, third touch = grey). **(B)** Peak firing rates distribution across 3 consecutive touches across all modulating electrodes. No significant difference between touches was observed for both participant C1 and P5. **(C)** Relationship between maximum z-scored firing rate and maximum applied force across trials for each digit in participant C1 during the *touch with vision* condition. No statistically significant correlation was found between force and firing rate.

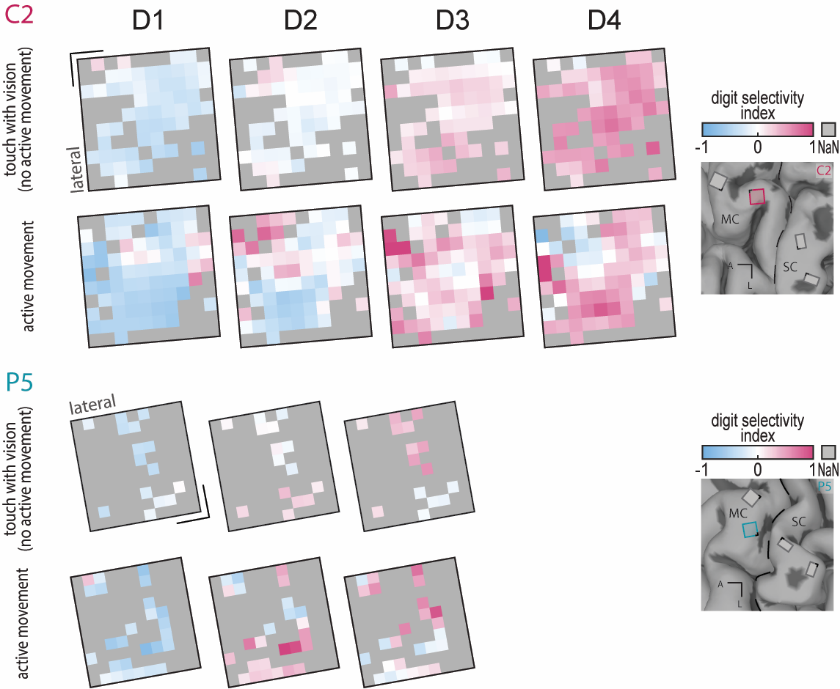

**Supp. Fig. 6. Touch-evoked responses in the human MC.** Maps of digit selectivity index for each electrode of the motor arrays during tactile stimulation of different digits (first and third row) and during attempted individual digits movements (second and forth row) in C2 and P5. Pink indicates higher neural activity for a given digit at a given electrode compared with the other digits recorded from the same electrode. Non-modulating electrodes are shown in grey. fMRI images for each participant with the related arrays’ placements are shown on the right. The arrays considered in this analysis are depicted in green or red.

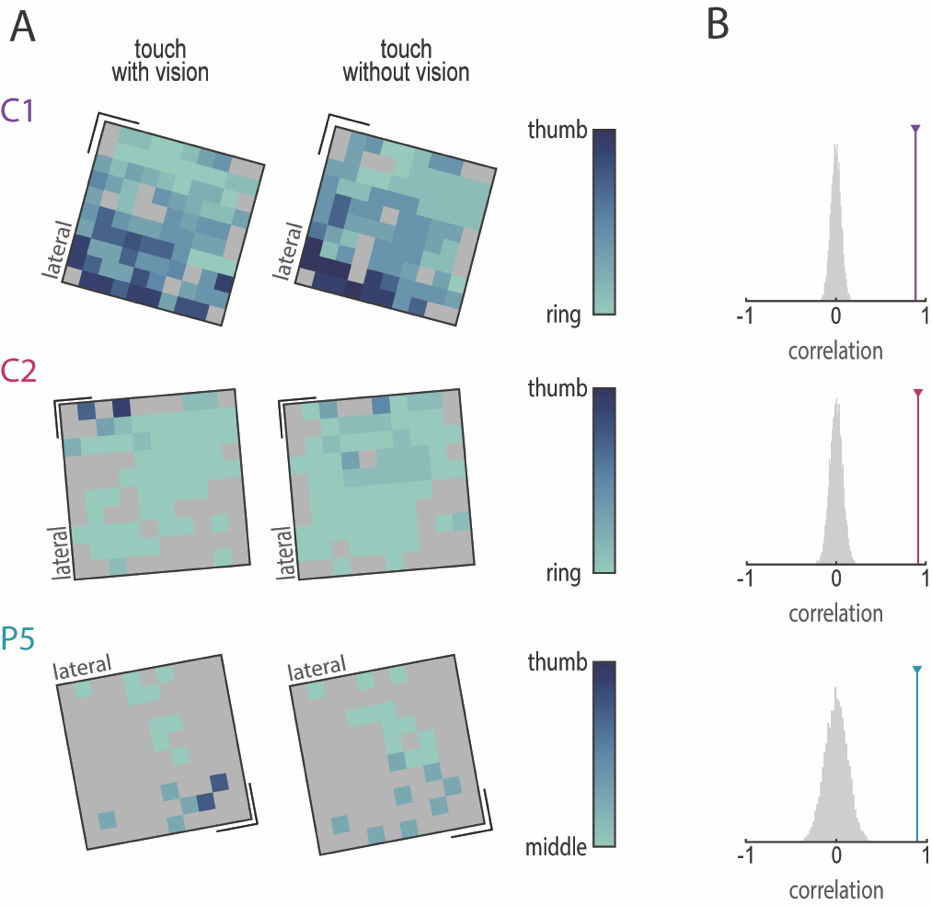

**Supp. Fig. 7. Touch-evoked response in absence of vision in MC.** **(A)** Digit selectivity gradient maps for each participant during touch with vision (first column) and without vision (second column). Dark blue indicates channels responding more strongly to tactile stimulation of the thumb and index finger, while light blue indicates electrodes responding more strongly to stimulation of the middle and ring fingers. **(B)** Correlation of digit preference between touch with vision and touch without vision conditions for each participant (colored line). The grey distribution shows the results obtained by recomputing the same correlation 10,000 times shuffling the digit selectivity index.

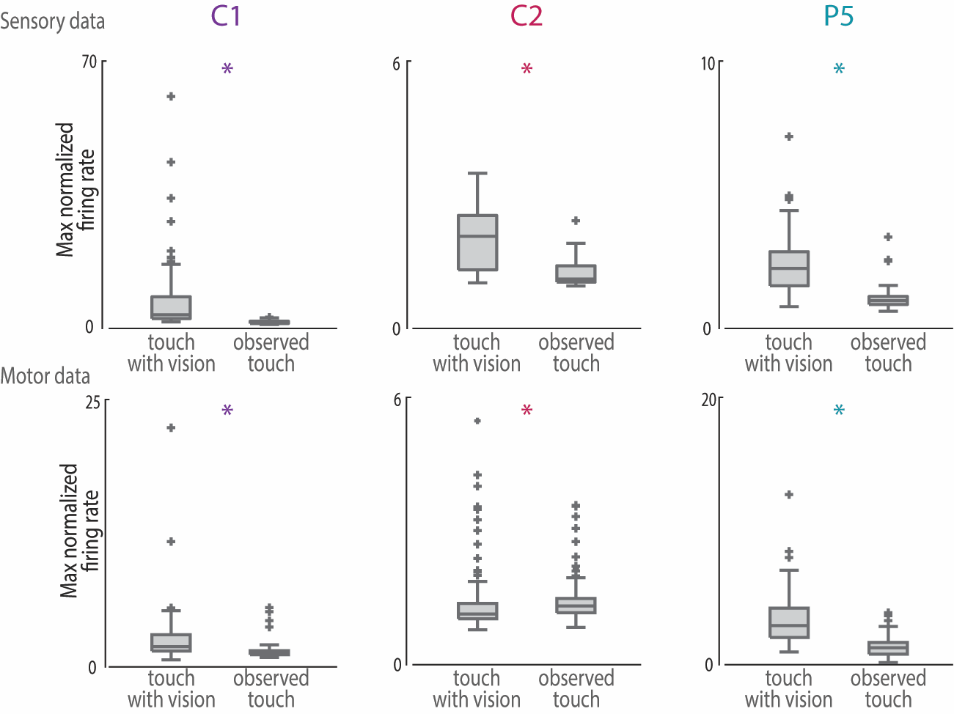

**Supp. Fig. 8. Maximum normalized firing rate distribution during touch with vision and observed touch.** Maximum normalized firing rate (firing rate/baseline) distribution during touch with vision and observed touch for the three participants (C1, C2, P5). Panels in the top row correspond to sensory-related neural activity, whereas panels in the bottom row correspond to motor-related neural activity. * indicates *p<0.05*.

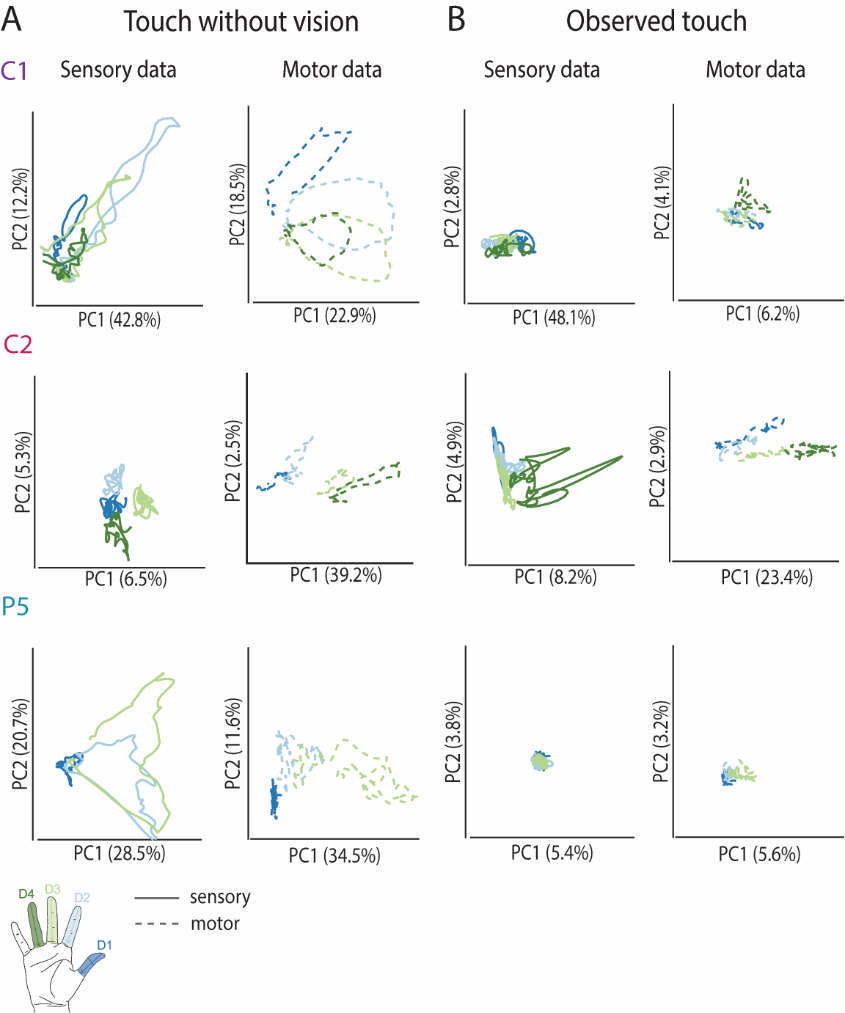

**Supp. Fig. 9. Cortical digits representation during touch. (A)** Neural activity from the sensory arrays (left) and motor arrays (right) during tactile stimulation without vision projected into a 2D PC space for the three participants. Each color represents one digit (D1-D4 for C1 and C2; D1-D3 for P5). **(B)** Neural activity from the sensory arrays (left) and motor arrays (right) during observed touch projected into a 2D PC space for the three participants. Each color represents one digit (D1-D4 for C1 and C2; D1-D3 for P5).

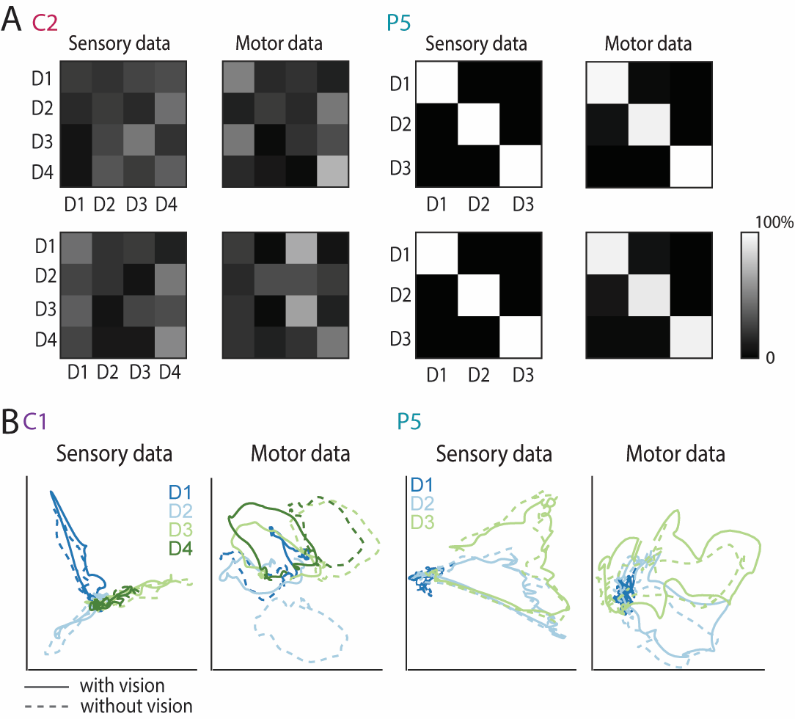

**Supp. Fig. 10. Cross-decoding of digits between touch with and without vision. (A)** Confusion matrices obtained using a model trained on the touch with vision condition and tested on the touch without vision condition (first row), and vice versa (second row) for C2 and P5 participants during touch window (D1-D4 for C2; D1-D3 for P5). **(B)** Neural activity from the sensory arrays (left) and motor arrays (right), computed from the combined dataset of touch with vision and touch without vision conditions, projected into a 2D PC space for C1 and P5. Each color represents one digit. The solid line represents the touch with vision condition, and the dotted line represents the touch without vision condition (D1-D4 for C1; D1-D3 for P5).

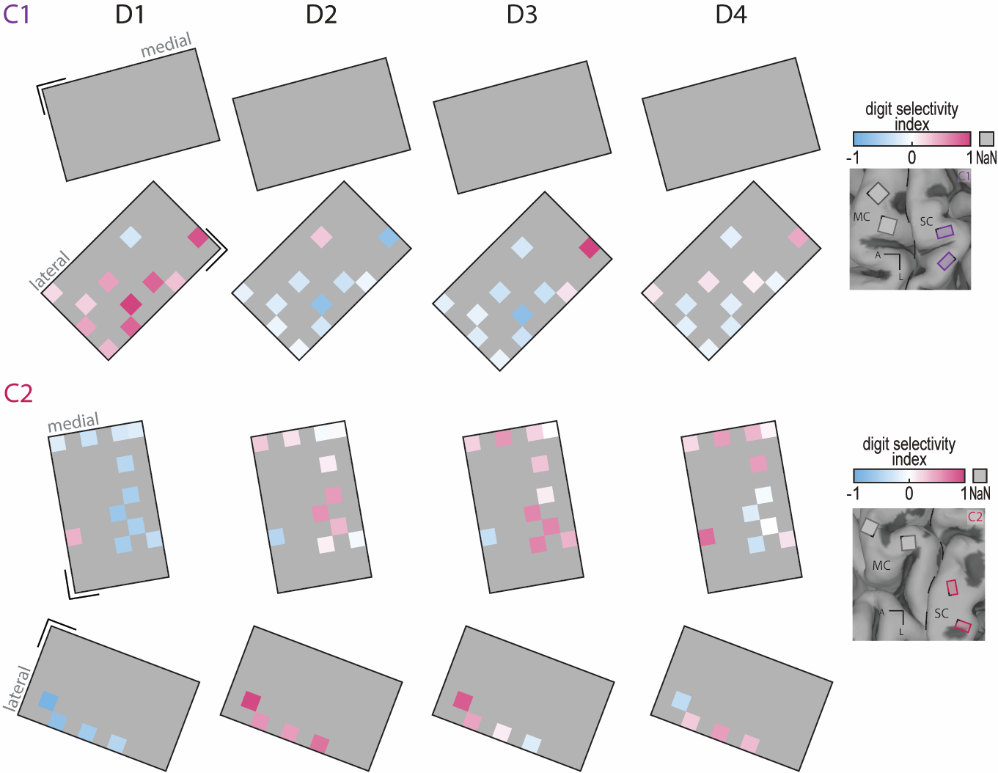

**Supp. Fig. 11. Attempted individual finger movements responses in the human SC.** Maps of the digit selectivity index for each electrode of the sensory arrays during attempted individual finger movements in C1 and C2. Pink indicates higher neural activity for a given digit at a given electrode compared with the other digits recorded from the same electrode. Non-modulating electrodes are shown in grey. MRI images for each participant with the related arrays’ placements are shown on the right. The arrays considered in this analysis are depicted in purple or red.

**Supp. Table S1.** **Tactile thresholds of the hand before implantation for each participant.**

|  | C1 | C2 | P5 |
| --- | --- | --- | --- |
| Mechanical sensory threshold at implant (g) |  |  |  |
| Thumb | 0.6 | >60 | 0.16 |
| Index | 0.6 | >60 | 0.4 |
| Middle | 2 | 60 | 0.6 |
| Ring | 1 | 26 | 2 |
| Pinky | 1 | 15 | 2 |
| Palm | 2 | - | 1.2 |
